# Targeting Lysosomal MCOLN1/TRPML1 Ion Channels to Alleviate Diabetes Mellitus via a Ca^2+^- CaMKKβ-AMPK pathway

**DOI:** 10.64898/2026.06.16.732532

**Authors:** Jiaxuan Zhu, Zhaonan Pan, Shijia Wang, Haoxiang Zhang, Qianhui Wang, Zikai Jiang, Yuyuan Wang, Zongxian Ding, Dan Li

## Abstract

Type 2 Diabetes mellitus (T2DM) is a metabolic syndrome characterized by hyperglycemia and various complications. Current drugs are limited by side effects and resistance, necessitating novel targets and therapies. Previous studies have shown that MK-83, a synthetic agonist of transient receptor potential mucolipin 1 (TRPML1/ MCOLN1), a lysosomal Ca^2+^ channel, activates adenosine 5′-monophosphate-activated protein kinase (AMPK), a key therapeutic target in diabetes. However, whether targeting TRPML1 can treat T2DM remains unclear. In this study, we found that transgenic overexpression or pharmacological activation of TRPML1 regulates AMPK phosphorylation via a Ca^2+^-CaMKKβ-dependent mechanism. This activation promotes glucose transporter 4 (GLUT4) translocation and increases intracellular glucose uptake. Conversely, genetic inactivation or pharmacologically inhibition of TRPML1 blocks both AMPK activation and glucose uptake. More importantly, pharmacological activation of TRPML1 *in vivo* alleviates hyperglycemia in db/db mice (a T2DM model), as evidenced by random blood glucose levels, fasting blood glucose levels and HbA1c levels. Furthermore, other hallmark features of db/db mice-including impaired oral glucose tolerance, reduced insulin tolerance, and elevated ALT and AST levels-were all ameliorated upon TRPML1 activation. Hence, targeting lysosomal TRPML1 channel represents a promising therapeutic strategy for T2DM, with highly specific TRPML1 agonists as potential anti-diabetic agents.

## Introduction

Type 2 diabetes mellitus (T2DM) is a chronic disease, characterized by hyperglycemia and relative insulin deficiency that affects more than 400 million people worldwide [1]. Despite the wide range of available antidiabetic drugs, side effects, drug resistance, and patient heterogeneity highlight the urgent need for new therapeutics with novel mechanisms to address clinical needs [2]. Eukaryotes possess a conserved signaling pathway that responds to low cellular ATP (due to reduced oxygen or glucose) by activating adenosine monophosphate-activated protein kinase (AMPK) [3], which then phosphorylates key targets to regulate lipid and glucose metabolism [4]. Given that AMPK is essential for glucose homeostasis [3], it represents a key therapeutic target for diabetes and related metabolic diseases. Mechanistically, AMPK is activated when its α-subunit is phosphorylated at Thr172 by upstream kinases via energy- or Ca²⁺-dependent pathways [3, 5]. Once activated, AMPK suppresses anabolic (energy-consuming) processes and promotes catabolic (energy-producing) ones [6]. In nutrient-rich conditions, ATP binds to the γ subunit of AMPK and keeps it inactive. Under energetic stress, AMP replaces ATP, triggering liver kinase B1 (LKB1) to phosphorylate the α subunit of AMPK.

Notably, cytoplasmic Ca^2+^ can activate AMPK independently of cellular energy status [5, 7]. Lysosomes, mitochondria and endoplasmic reticulum (ER) are the main intracellular Ca^2+^ stores [8] and some connections between distinct cytosolic Ca^2+^ sources and AMPK activation have been investigated [9]. Nevertheless, the involvement of lysosomes in AMPK activation and its implications for treating diabetes and metabolic diseases have yet to be elucidated. Transient Receptor Potential Mucolipin 1 (MCOLN1/TRPML1) is the best characterized lysosomal Ca^2+^ channel, mediating Ca^2+^ release from the lumen to the cytosol. TRPML1 dysfunction underlies various metabolic diseases, lysosomal storage diseases and neurodegenerative disorders [10–12]. For example, mutations in the gene encoding TRPML1 cause mucolipidosis type IV (MLIV), a lysosomal storage disease characterized by early-onset progressive neurodegeneration [13]. Lysosomal dysfunction is closely associated with multiple metabolic diseases, including obesity, T2DM, non-alcoholic fatty liver disease, and atherosclerosis [14, 15]. Impaired lysosomal function disrupts lipid metabolism, autophagy, and inflammation, leading to insulin resistance and β-cell failure [16].

Previous studies have shown that the synthetic TRPML1 agonist MK6-83 activates AMPK in human fibroblasts [9]. However, whether the TRPML1–AMPK axis regulates glucose metabolism, particularly in the context of T2DM, remains unclear. To address this, the current study investigates how TRPML1 activation regulates glucose metabolism, and ML-SA8, a TRPML1 agonist with higher *in vitro* potency (ML-SA8>ML-SA5 > MK6-83) [17–19], was applied to elucidate the therapeutic effects and molecular mechanisms of TRPML1 activation in T2DM models.

In the current proof-of-concept study, we found that both genetic and pharmacological activation of the lysosomal TRPML1 channel are sufficient to activate AMPK and alleviate diabetic phenotypes *in vitro* and *in vivo*. Therefore, we propose that pharmacological activation of TRPML1 may serve as a potential therapeutic strategy for T2DM.

## Results

### Pharmacological activation of lysosomal TRPML1 induces AMPK activation

Cytosolic Ca^2+^ reportedly activates AMPK [20, 21]. Lysosome, mitochondria and ER are the major compartmentalized Ca^2+^ stores in cells [8]; the mechanistic links between cytosolic Ca^2+^ and AMPK activation lead us to consider the discrete Ca^2+^ sources as a potential specificity determinant of AMPK activation [22, 23]. In this study, we investigated whether lysosome-released ions can activate AMPK, whose activation was measured by the phosphorylation status of threonine 172 (referred as Thr172) [24].

TRPML1 is the principal Ca^2+^-permeable channel that releases Ca^2+^ from the lumen into the cytosol upon cellular stimulation [25, 26].Thus we then investigated whether TRPML1 contributes to AMPK activation. Although the synthetic TRPML1 agonist MK6-83 was previously shown to activates AMPK in human fibroblasts [9], here we used a much more potent TRPML1 agonist, ML-SA8 [27] (EC_50_=0.086 ± 0.013 μM for endogenous TRPML1; ***Suppl. Fig. 1a, b***). As shown in **Fig. 1a, b**, ML-SA8 (0.3-3 μM, 2 h) induces robust AMPK phosphorylation in a dose-dependent manner in HepG2 cells as well as several other cell lines including HeLa (***Suppl. Fig. 1c, d***) and HAP1 (***Suppl. Fig. 1e, f***). And this activation by ML-SA8 was completely blocked by dorsomorphin/compound C (CC, 10 μM, an AMPK inhibitor [28]) (**Fig. 1c, d**). Similarly, AMPK can be activated by another less potent TRPML1 agonist ML-SA5 (potency SA8> SA5>MK6-83 [18, 27, 29]) in HepG2, HeLa and HAP1 cells (***Suppl. Fig. 2a-c)***. ML-SA8 (1 μM) increased the phosphorylation of AMPK was further confirmed by increased phosphorylation of acetyl-CoA carboxylase (ACC-Ser79), a direct substrate of AMPK (***Suppl. Fig. 2d***). Moreover, ADP/ATP ratios remained not affected with ML-SA8 treatment (***Suppl. Fig. 2e***), indicating that ML-SA8-induced AMPK activation was independent of energy stress. Note that the specificities of ML-SAs have been previously validated using TRPML1 knockout (KO) cells [18, 29] and confirmed in the atomic-resolution co-structures [30, 31]. In addition, we investigated the role of another lysosomal Ca²⁺ channel, TPC2, using its specific agonist riluzole [25]. As shown in ***Suppl. Fig. 3a, b***; AMPK activation required 800 μM of riluzole, a concentration exceeding its physiological range [32–35]. Therefore, we focused our research on ML-SA8-induced AMPK phosphorylation.

**Figure 1.**
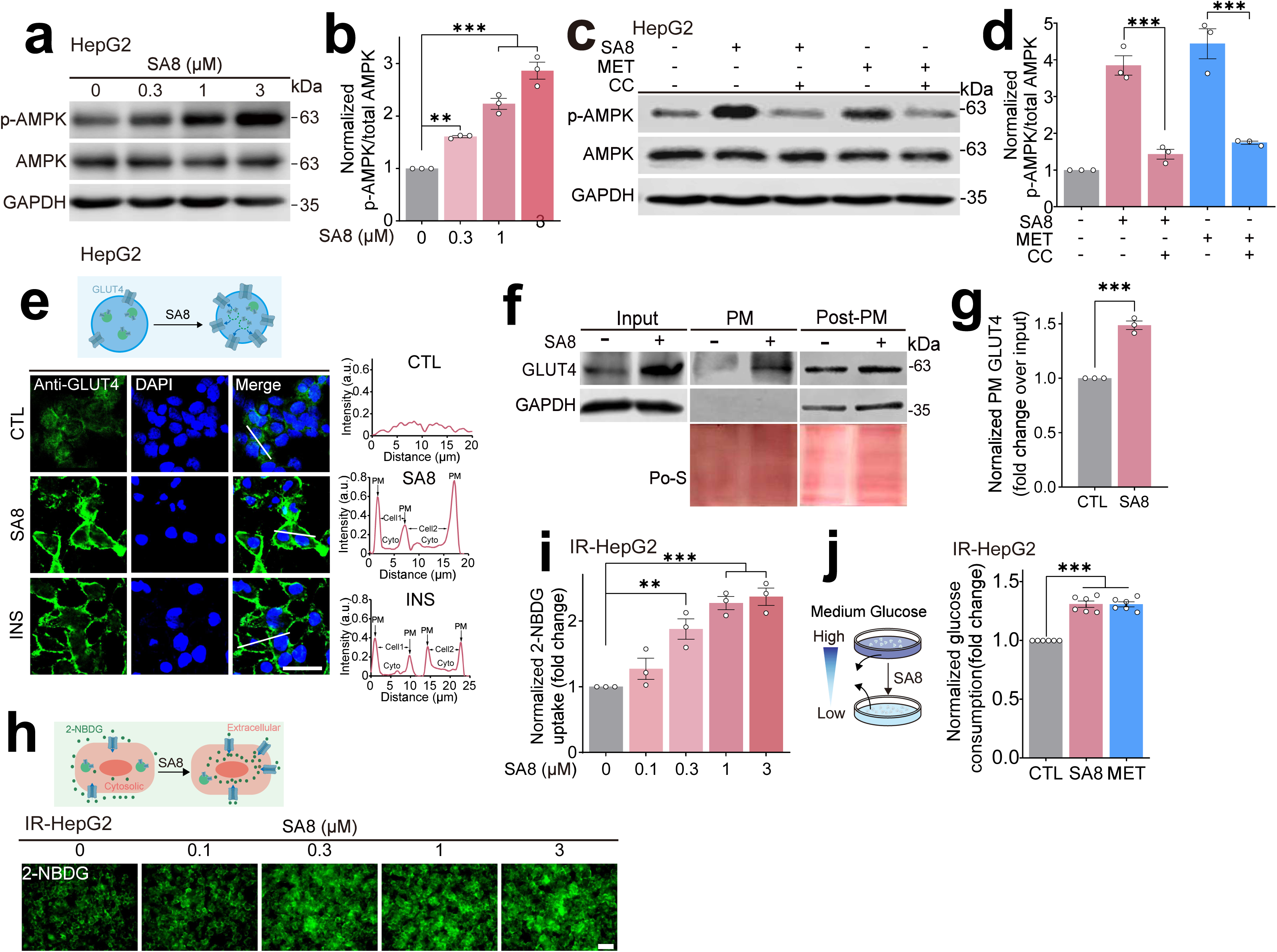
Stimulation of the lysosomal TRPML1 channel induces AMPK activation. (**a**) ML-SA8 (a TRPML1 specific synthetic agonist) activated AMPK in a dose-dependent manner. HepG2 cells were treated with ML-SA8 (0.3-3 μM) for 2 h. (**b**) Quantification of results shown in **c.** from n= 3 independent experiments. **(c)** AMPK inhibitor blocked ML-SA8-induced AMPK phosphorylation. HepG2 cells were pretreated with compound C (CC, 10 μM, 1h), followed by cotreatment with ML-SA8 (1 μM)/ Metformin (MET, 2 mM) for 2 h and AMPK phosphorylation was measured by Western blotting. **(d)** Ratio of p-AMPK/total AMPK shown in **e.** (n= 3). (**e**) ML-SA8 induced GLUT4 PM translocation. HepG2 cells were treated with ML-SA8 (1 μM, 4 h) and GLUT4 immunoreactivity was measured. Nuclei were counterstained with DAPI (blue). Scale bar, 20 μm. The graph at the *Righter* panel shows the fluorescence intensity of GLUT4 with a white line scan across a cell (white line shown in the *Lefter* images). (**f**) Western blot analysis of PM GLUT4 expression induced by ML-SA8. Membrane and cytoplasm protein fractions were isolated from cells treated with ML-SA8 (1 µM) or DMSO for 4 h and subjected to western blot analysis. GAPDH served as the loading control for the Input and Post-PM fractions. Ponceau S (Po-S) staining was used as the loading control for the PM fraction. (**g**) Quantification of PM GLUT4 expression normalized to input as shown in **f**. (n= 3 independent experiments). (**h**) ML-SA8 promoted glucose influx. IR-HepG2 cells were treated with ML-SA8 (0.1-3 μM) for 4 h, followed by 2-NBDG (50 µM, 0.5 h) treatment. Scale bar, 40 μm. (**i**) Quantification of the relative fluorescent intensity of 2-NBDG in **h.** from n= 3 independent experiments. (**j**) The effect of ML-SA8 on glucose consumption in medium. IR-HepG2 cells were treated with or without ML-SA8 (1 μM) for 4 h in DMEM containing 10 mM glucose, no FBS, no phenol red, and glucose consumption in the medium was measured. MET (2 mM) served as a positive control. For all panels, data are presented as mean ± s.e.m.; \**p* < 0.05, \*\**p* < 0.01, \*\*\**p* < 0.001, ANOVA.

Glucose transporters (GLUTs) at the plasma membrane reportedly facilitate glucose uptake by an inward diffusion gradient for glucose [36]. In particular, GLUT4 is known to participate in AMPK-mediated glucose uptake by translocating from cytoplasm to plasma membrane [37]. Although GLUT4 is not the predominant GLUT isoform in the liver, it is nevertheless expressed in this organ [38–40]. Therefore, we next investigated whether ML-SA8 regulates the translocation of GLUT4. As shown in **Fig. 1e**, ML-SA8 (1 μM) induced a substantial increase in GLUT4 protein at the plasma membrane (PM) of HepG2 cells by immunofluorescence, indicating that ML-SA8 promotes GLUT4 translocation to the PM. Moreover, we isolated PM and post-PM fractions (supernatants devoid of PM) from HepG2 cells treated with ML-SA8 or vehicle control. Quantification of GLUT4 levels in the PM fraction confirmed that ML-SA8 stimulated GLUT4 translocation (**Fig. 1f, g**). Moreover, ML-ML-SA8 upregulated both the mRNA and expression level of GLUT4 in HepG2 cells (***Suppl. Fig. 4a***, **Fig. 1f**, ***Suppl. Fig. 4b***). Collectively, these finding indicate that ML-SA8 promotes both PM translocation and the expression of GLUT4.

We then studied the effect of ML-SA8 on glucose levels by measuring both cellular glucose uptake and glucose consumption in the medium [41]. First, we constructed a palmitic acid (PA)-induced insulin resistance (IR) cell model [42, 43]. As shown in ***Suppl. Fig. 5a, b***, insulin (100 nM, 0.5 h) induced glucose influx in HepG2 cells measured by 2-deoxy-2-[(7-nitro-2,1,3-benzoxadiazol-4-yl)amino]-d-glucose (2-NBDG), a fluorescent glucose analog, which is transported into cells and phosphorylated by hexokinase, trapping it intracellularly and allowing fluorescence-based quantification of glucose uptake [44]. In contrast, in PA (0.25 mM, 12 h)-treated HepG2 cells, insulin (100 nM, 0.5 h)-induced glucose influx was almost completely blocked, suggesting that PA-treated HepG2 cells are less sensitive to insulin (Thereafter IR-HepG2 represents PA-induced insulin resistant HepG2 cells). ML-SA8 (1 μM, 2 h)-induced AMPK phosphorylation was validated in IR-HepG2 cells by western blot analysis (***Suppl. Fig. 5c***). In IR-HepG2 cells, we observed that upon ML-SA8 (0.1-3 μM, 2-6 h) treatment, the intensities of 2-NBDG were dramatically increased compared to control in a dose-(**Fig. 1h, i)** and time-dependent (***Suppl. Fig. 5d, e***) manner. Correspondingly, glucose consumption in the culture medium was dramatically decreased with ML-SA8 (1 μM, 4 h) treatment in IR-HepG2 cells (**Fig. 1j**). Likewise, ML-SA8 (1 μM) was sufficient to induce cellular glucose uptake in various cells such as human hepatocellular carcinoma cells (Huh7) and mice myoblast cells (C2C12) following IR (PA-treated) induction (***Suppl. Fig. 5f, g***). Taken together, these data demonstrated that stimulation of TRPML1 activates AMPK signaling cascades, which facilitates GLUT4 PM translocation, leading to the increases of cellular glucose uptake.

### TRPML1 is essential in ML-SA-induced AMPK activation

We then investigated whether increasing the expression of TRPML1 in HepG2 cells is sufficient to increase the cells’ sensitivity to ML-SA8 treatment. Remarkably, a low dose of ML-SA8 (0.03 μM) can activate AMPK in HepG2 cells overexpressing GFP-TRPML1, but not in wild type (WT) cells (**Fig. 2a**). Consistently, ML-SA8 (0.03 μM) can also promote cellular glucose uptake in TRPML1-overexpressing IR-HepG2 cells (**Fig. 2b, c**), while no such effects were observed in IR-HepG2 cells. Collectively, these data demonstrate that the TRPML1 overexpression can promote AMPK activation.

**Figure 2.**
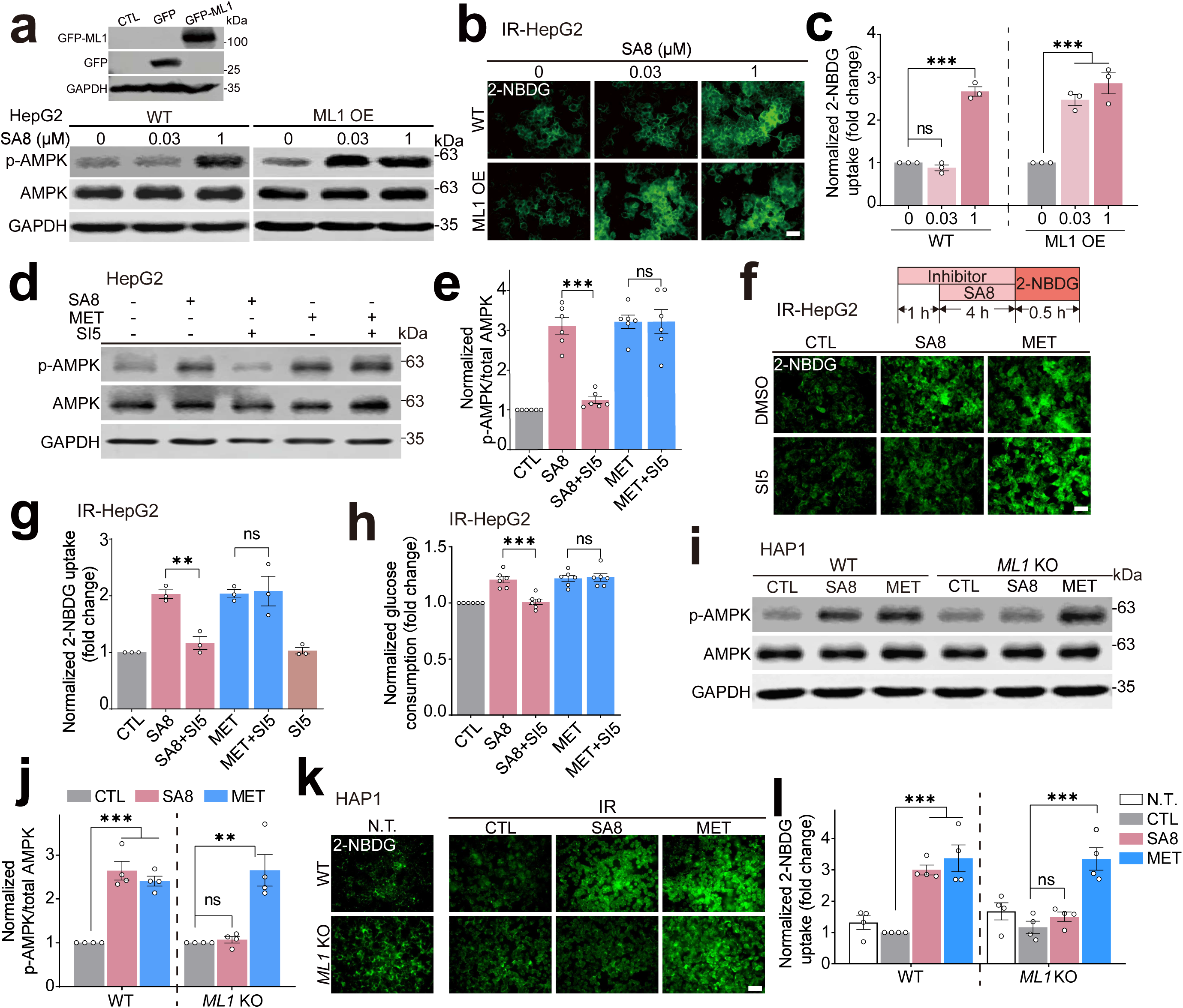
TRPML1 is required for ML-SA8-induced AMPK activation. **(a)** Overexpressing *TRPML1* further promoted low-dose of ML-SA8-activating AMPK. In the top panel, western blot validation of the GFP-TRPML1 overexpression in HepG2. In the bottom panel, HepG2 cells with or without GFP-TRPML1 overexpression were treated with ML-SA8 (0.03-1 µM) for 2 h and western blotting analysis of p-AMPK/total AMPK protein levels. **(b)** TRPML1 overexpression promoted cellular glucose uptake (2-NBDG) by low-dose of SA8 (0.03-1 µM, 4 h) in IR-HepG2 cells. Scale bar, 40 μm. **(c)** Quantification of the relative fluorescent intensity of 2-NBDG shown in **b.** (n= 3 independent experiments). (**d**) ML-SI5 (a TRPML1 synthetic inhibitor) blocked ML-SA8-induced AMPK activation. Western blot analysis of AMPK phosphorylation in HepG2 cells pretreated with ML-SI5 (3 μM) for 1 h, followed by cotreated with ML-SA8 (1 μM) or MET (2 mM) for 2 h. (**e**) Quantification of the results shown in **d.** from n= 6 independent experiments. (**f**) The effect of ML-SI5 on ML-SA8-induced cellular glucose uptake. IR-HepG2 were pretreated with ML-SI5 (3 μM) for 1 h, then cotreated with ML-SA8 (1 μM, 4 h), followed by 2-NBDG (50 µM, 0.5 h). Scale bar, 40 μm. (**g**) Quantification of the intensity of 2-NBDG in **f.** (n= 3). (**h**) ML-SI5 blocked ML-SA8-induced glucose consumption. IR-HepG2 cells were pretreated with ML-SI5 (3 μM) for 1 h, then cotreated with ML-SA8 (1 μM) or MET (2 mM) for 4 h, finally glucose levels in the supernatant medium were measured. (**i**) Western blotting analysis of p-AMPK levels in WT and *TRPML1* KO HAP1 cells treated with ML-SA8 (1 μM) or MET (2 mM) for 2 h. (**j**) Quantitative analysis of p-AMPK/total AMPK levels as shown in **i.** from n= 4 independent experiments. (**k**) *TRPML1* KO abrogated ML-SA8-but not MET-induced cellular glucose uptake. WT and *TRPML1* KO HAP1 cells were treated with PA (0.25 mM, 12 h), followed by ML-SA8 (1 μM) or MET (2 mM) for 4 h. 2-NBDG (50 µM, 0.5 h) was then applied and measured. Scale bar, 40 μm. (**l**) Quantification of the relative fluorescent intensity of 2-NBDG in **k.** from n= 4 independent experiments. In all panels, data are presented as mean ± s.e.m.; \*\**p* < 0.01, \*\*\**p* < 0.001.

Next, we investigated whether TRPML1 is required for ML-SA-induced AMPK activation using the TRPML1 specific synthetic inhibitor-ML-SI5, whose specificities have been previously validated [17, 27]. Pretreatment with ML-SI5 (3 μM) completely abolished ML-SA8-triggered (1 μM, 2 h) AMPK phosphorylation, while no effect on metformin (MET)-induced AMPK activation (**Fig. 2d, e**). Moreover, in IR-HepG2 model, ML-SI5 substantially inhibited ML-SA8-, but not MET-induced cellular glucose uptake and extracellular glucose consumption (**Fig. 2f-h**). To further validate the TRPML1-dependent mechanism, we utilized a *TRPML1* knockout (KO) HAP1 cell line constructed by CRISPR-Cas9 as previously described [45] and validated in ***Suppl. Fig. 6a***. In *TRPML1* KO HAP1 cells, ML-SA8 (1 μM, 2 h)-induced AMPK phosphorylation and cellular glucose uptake were almost completely abolished compared to wild-type (WT) HAP1 cells (**Fig. 2i-l**). Collectively, these data demonstrate that TRPML1 is both sufficient and necessary for ML-SA-induced AMPK activation in IR model cells.

### Ca^2+^ -CaMKKβ -dependence of ML-SA-induced AMPK activation

Lysosomal TRPML1 is a non-selective cation channel that is permeable to Ca^2+^, as well as heavy metal ions such as Fe^2+^ and Zn^2+^ [46]. Given the well-established role of Ca^2+^ in AMPK activation [47], we first examined whether Ca^2+^ is required for ML-SA’s AMPK activation upon TRPML1 stimulation. BAPTA-AM (20 μM), a membrane-permeable Ca^2+^ chelator (fast-acting) [48] readily blocked ML-SA8-evoked the phosphorylation of AMPK and its substrate ACC (**Fig. 3a, b, *Suppl. Fig. 6b***) and cellular glucose uptake (**Fig. 3c, d**). In contrast, EGTA-AM, another Ca^2+^ chelator (slow-acting), failed to block ML-SA8’s effects (**Fig. 3a-d**). The inhibitory effects of BAPTA-AM (20 μM) and EGTA-AM (20 μM) on cytosolic Ca^2+^ was confirmed in HEK293 cells overexpressing GCaMP7-TRPML1 (***Suppl. Fig. 6c***). Considering that BAPTA has much faster Ca^2+^ association and dissociation kinetics than EGTA [49], this disparate Ca^2+^-dependent mechanism suggests that ML-SA8-regulated AMPK cascades through a fast and local Ca^2+^ signal from lysosomes. Next, we examined whether Fe^2+^ and Zn^2+^ involve in ML-SA-triggered AMPK cascades. TPEN (10 μM), a high-affinity Zn²⁺ chelator [50], and deferoxamine (DFO) (10 μM), a Fe^2+^ chelator [51], both failed to block ML-SA8-induced AMPK activation and cellular glucose uptake (***Suppl. Fig. 6d-g***). Together, these data suggest that TRPML1 activation regulates AMPK-GLUT4-glucose pathway via a fast Ca^2+^ release from lysosomes.

**Figure 3.**
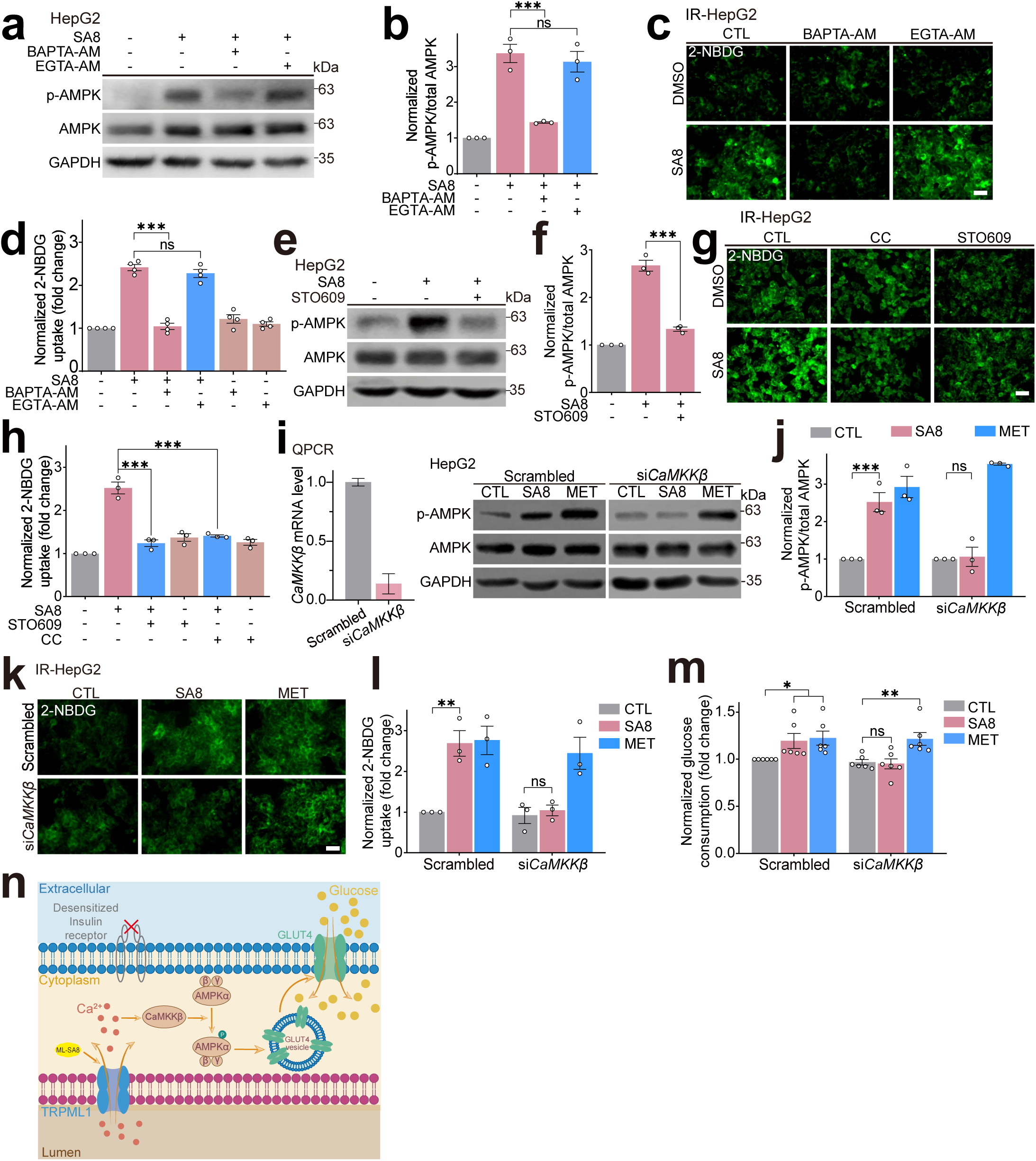
Ca^2+^- CaMKKβ dependence of ML-SA8 induced-AMPK activation. **(a)** BAPTA-AM inhibited ML-SA8-induced AMPK phosphorylation. HepG2 cells were pretreated with BAPTA-AM (fast Ca^2+^ chelator, 20 μM) or EGTA-AM (slow Ca^2+^ chelator, 20 μM) for 1 h, then cotreated with ML-SA8 (1 μM, 2 h), AMPK phosphorylation was then evaluated by Western blotting. **(b)** Quantification of results shown in **a.** from n= 3 experiments. **(c)** The effect of BAPTA-AM on ML-SA8-induced cellular glucose uptake. IR-HepG2 cells were treated with ML-SA8 (1 μM, 4 h) in the presence or absence of BAPTA-AM (20 μM) or EGTA-AM (20 μM), followed by analysis of cellular glucose levels with 2-NBDG. Scale bar, 40 μm. **(d)** Quantification of results shown in **c.** from (n= 4). **(e)** CaMKKβ inhibitor blocked ML-SA8-induced AMPK phosphorylation. HepG2 cells were pretreated with STO609 (10 μM, 1h) and cotreated with ML-SA8 (1 μM, 2 h), and AMPK phosphorylation was assayed by Western blotting. **(f)** Ratio of p-AMPK/total AMPK shown in **e.** (n= 3). **(g)** CC and STO609 abolished ML-SA8-induced cellular glucose uptake. IR-HepG2 cells were treated with ML-SA8 (1 μM) with or without CC (10 μM) or STO609 (10 μM) for 4 h and cellular glucose levels were measured by 2-NBDG. Scale bar, 40 μm. **(h)** Quantification of the intensity of 2-NBDG shown in **g.** (n= 3). **(i)** CaMKKβ KD abolished ML-SA8-induced AMPK activation. CaMKKβ KD efficiency was examined by QPCR (*Left panel*). HepG2 cells were transfected with CaMKKβ siRNA and then treated with ML-SA8 (1 μM) or MET (2 mM) for 2 h, then p-AMPK levels were measured by Western blot. **(j)** Quantitation of results shown in **i**. **(k)** CaMKKβ KD abrogated ML-SA8-induced cellular glucose uptake. IR-HepG2 cells were treated with ML-SA8 (1 μM) or MET (2 mM) for 4 h. 2-NBDG (50 µM, 0.5 h) was then applied and measured. Scale bar, 40 μm. **(l)** Quantification of results in **k**. (n= 3) **(m)** The effects of CaMKKβ KD on ML-SA8-induced glucose consumption. IR-HepG2 cells were treated with ML-SA8 (1 μM) or MET (2 mM) for 4 h, then glucose levels in the supernatant medium were measured (n=3). **(n)** A working model to illustrate that small-molecule TRPML1 agonist-ML-SA8 promotes cellular glucose uptake via TRPML1-Ca²⁺-CaMKKβ-AMPK-GLUT4 pathway. ML-SA8 specifically activates TRPML1 channel, which releases Ca²⁺ into the cytoplasm. Elevated cytosolic Ca²⁺ then triggers CaMKKβ kinase activation, which phosphorylates AMPK to initiate its catalytic activity, followed by promoting GLUT4-containing vesicle translocation to the PM. Membrane-inserted GLUT4 facilitates extracellular glucose uptake into cells, ultimately restoring glycemic homeostasis. Arrows denote sequential signaling events, with color-coded molecular entities matching experimental annotations.

We next investigated how ML-SA8 induces AMPK phosphorylation. A key mechanism of AMPK activation involves phosphorylation of its subunit at Thr172 site by upstream kinases, including CaMKKβ (calcium/calmodulin-dependent protein kinase kinase β which responds to elevated cytosolic Ca^2+^ [52, 53] and LKB1 [7]. Considering our finding that TRPML1-dependent Ca^2+^ release is essential for AMPK activation (**Fig. 3a-d**), we hypothesize that CaMKKβ serves as the critical upstream kinase bridging lysosomal Ca²⁺ signaling and ML-SA8-stimulated AMPK activation. Notably, STO609 (a CaMKKβ inhibitor, 10 μM) [54] significantly suppressed ML-SA8-induced AMPK phosphorylation in HepG2 cells (**Fig. 3e, f**). Furthermore, the effects of ML-SA8 on cellular glucose uptake (2-NBDG) were completely blocked by STO609 or CC (**Fig. 3g, h**). To further validate these findings, a genetic knockdown approach was employed. Consistent with the pharmacological inhibition, CaMKKβ knockdown markedly attenuated ML-SA8-induced AMPK phosphorylation **(Fig. 3i, j)** and abolished the associated increases in glucose uptake and extracellular glucose consumption **(Fig. 3k–m)**. Collectively, these findings establish CaMKKβ as an essential upstream mediator required for AMPK activation and the improved glucose metabolism following ML-SA8 treatment.

Overall, these *in vitro* experiments demonstrate that pharmacological activation of TRPML1 stimulates AMPK via a Ca^2+^-CaMKKβ-dependent pathway, thereby promoting GLUT4 translocation to the PM. This process enhances intracellular glucose uptake and helps maintain glucose homeostasis in a hepatic IR model (**Fig. 3n**).

### TFEB-independence of ML-SA-induced AMPK activation

TRPML1 activation is known to upregulate lysosomal biogenesis through TFEB activation [48, 55, 56]. While AMPK reportedly mediates phosphorylation of TFEB, and its transcriptional activity [57]. TFEB belongs to the microphthalmia (MiT/TFE) family, which also contains TFE3 and MITF, contributing to autophagic and lysosomal biogenesis [55, 58]. Hence, we examined whether TFEB/TFE3/MITF participates in TRPML1 stimulation-induced AMPK activation by using *TFEB/TFE3/MITF* triple KO (TKO) cells. The efficiency of *TFEB/TFE3/MITF* TKO was confirmed by Western blotting with anti-TFEB, TFE3 and MITF specific antibodies (***Suppl. Fig. 7a***). We found that in TKO cells, upon ML-SA8 (1 μM, 2 h) treatment, the increase of intensity of p-AMPK/total AMPK was not affected (***Suppl. Fig. 7b, c***), suggestive of TFEB/TFE3/MITF-independence of ML-SA8-triggered AMPK activation.

Next, we also assessed the role of AMPK in TRPML1 activation-mediated TFEB activity. Consistent with previous studies [18], ML-SA8 treatment (1 μM, 4 h) led to GFP-TFEB translocation from cytosol to nuclei in HeLa GFP-TFEB stable cells (***Suppl. Fig. 7d, e***), and this effect can be blocked by ML-SI5 treatment. However, CC or STO609 treatment failed to block ML-SA8-induced TFEB nuclear translocation (***Suppl. Fig. 7d, e***), suggesting that AMPK is dispensable for TRPML1 activation-induced subcellular TFEB translocation. Taken together, these results suggest that ML-SA8 -induced TFEB nuclear translocation is AMPK-independent.

### Administration of ML-SA8 ameliorates diabetic phenotypes in db/db mice

Hepatic steatosis is strongly associated with T2DM and is considered as an exacerbating factor of hepatic insulin resistance. We next investigated whether the small-molecule TRPML1 agonist ML-SA8 could alleviate diabetic phenotypes in C57BLKS/J-Lepr KO (*db/db*) mice, a well-established diabetic mouse model [59]. To determine whether ML-SA8 targets/activates AMPK *in vivo*, 9-week-old *db/db* mice were intraperitoneally (i.p.) injected with ML-SA8 (4 mg/kg) or vehicle (5% DMSO, 90% PEG300, 5% ddH_2_O). After 24 h, liver tissues were collected and pT172-AMPK/AMPK levels were measured by western blotting. As shown in **Fig. 4a, b**, ML-SA8 treatment significantly increased pT172-AMPK protein levels in liver tissues compared to vehicle, demonstrating that ML-SA8 activates AMPK in the liver of *db/db* mice.

**Figure 4.**
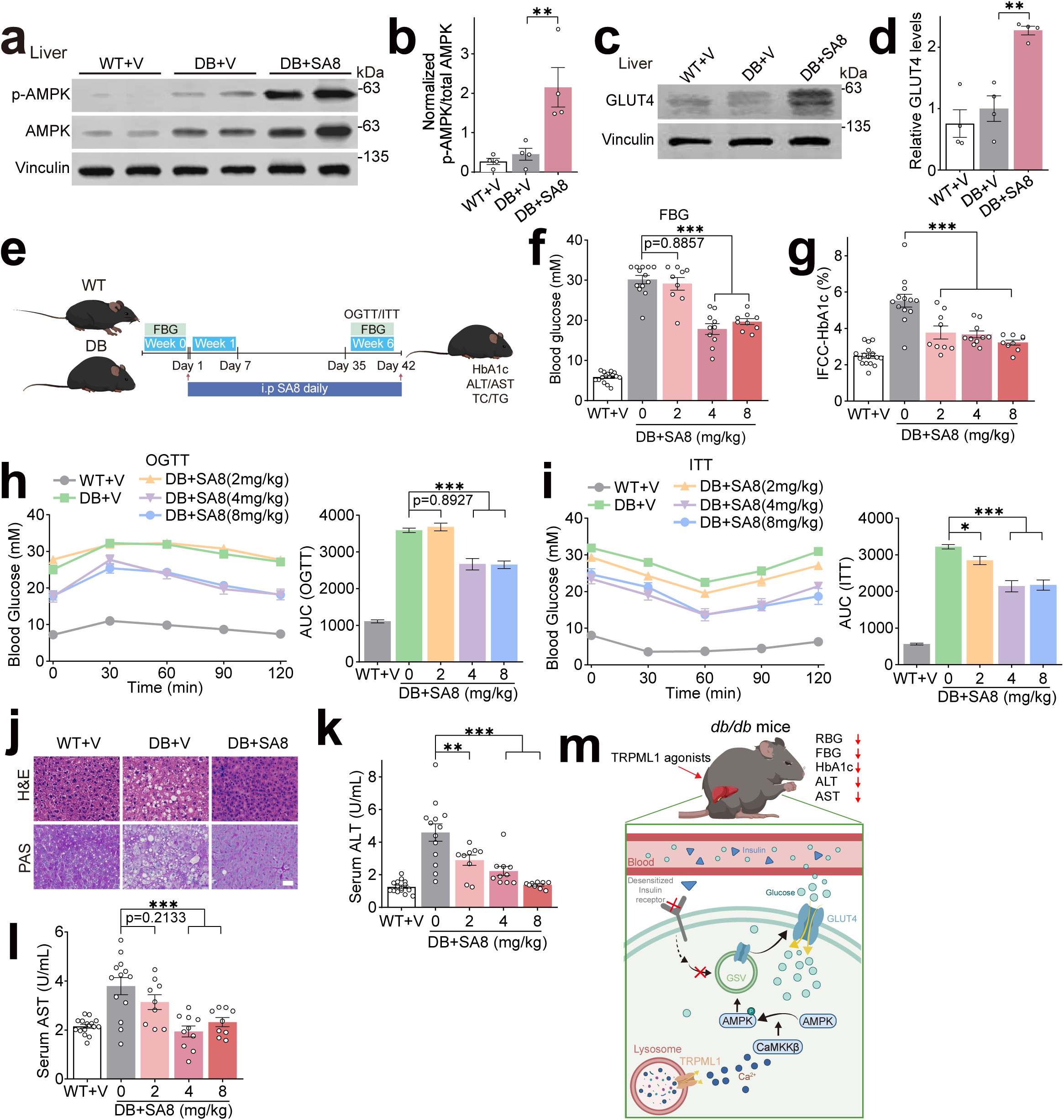
Administration of ML-SA8 attenuates metabolic disorder in *db/db* mice. **(a)** ML-SA8 promoted AMPK phosphorylation in the liver tissues of *db/db* mice with ML-SA8 (4 mg/kg) or vehicle (5% DMSO, 90% PEG300, 5% ddH_2_O) for 24 h. Liver tissues were collected and extracted proteins were subjected to detect p-AMPK/total AMPK by Western blotting. **(b)** Quantification of results shown in **a**. (n= 4). **(c)** ML-SA8 promoted GLUT4 expression in the liver tissues of *db/db* mice with ML-SA8 (4 mg/kg) or vehicle (5% DMSO, 90% PEG300, 5% ddH_2_O) for 6 weeks. Liver tissues were collected and extracted proteins were subjected to detect GLUT4 proteins by Western blotting. **(d)** Quantification of results shown in **c**. (n= 4). **(e)** Experimental design. *db/db* mice were randomly allocated into 4 groups: DB+V group was i.p. injected with vehicle (5% DMSO, 90% PEG300, 5% ddH_2_O) daily (n=11); DB+SA8 group was i.p. injected with SA8 (2mg/kg, n=9; 4mg/kg, n=10; 8 mg/kg, n=9). **(f)** Fasting blood glucose (FBG) levels at 0 and 6 weeks. **(g)** HbA1c levels. **(h)** OGTT. Mice daily i.p with SA8 (2, 4, 8 mg/kg) received oral glucose (1g/kg) after 10 h fasting, followed by blood glucose dynamic measurement (0-120 min) with corresponding AUC at the *Right* panel. **(i)** ITT. Mice with SA8 (2, 4, 8 mg/kg) treatment were i.p injected insulin (0.75 U/kg) after 4 h fasting, followed by blood glucose measurements. Left panel: Blood glucose levels (0-120 min). Right panel: AUC values. **(j)** Representative photomicrographs of H&E and PAS staining in liver tissues from mice treated with SA8 (4 mg/kg). Scale bar, 40 μm. **(k)** Serum ALT levels. **(l)** Serum AST levels. All data are presented as mean ± s.e.m.; \**p* < 0.05, \*\**p* < 0.01, \*\*\**p* < 0.001, ANOVA. **(m)** Pharmacological up-regulation of TRPML1 ameliorates diabetic phenotypes *in vivo* through Ca^2+^-dependent AMPK activation. ML-SA8 (a specific TRPML1 agonist) activates lysosomal TRPML1 channel, leading to lysosomal Ca^2+^ release from lysosomes. Ca^2+^ triggers Ca^2+^-dependent AMPK phosphorylation and initiates GLUT4 Storage Vesicle (GSV) trafficking to the PM. Subsequently, extracellular glucose is transferred into cells via GLUT4 and blood glucose levels are reduced *in vitro* and *in vivo*.

We next evaluated the *in vivo* therapeutic efficacy of ML-SA8. 9-week-old *db/db* mice were randomly divided into three groups and treated daily for 6 weeks: 1) Vehicle group (DB+V): i.p. injection of vehicle (5% DMSO, 90% PEG300, 5 % ddH_2_O); 2) ML-SA8 group (DB+SA8): i.p. injection of ML-SA8 at 2, 4, 8 mg/kg. We further validated the targeting of AMPK in various tissues by 6-week treatment of ML-SA8. ML-SA8 induced AMPK phosphorylation levels in liver (***Suppl. Fig.8a***), as well as in adipose (***Suppl. Fig.8b***) and skeletal muscle tissues (***Suppl. Fig.8c***) in *db/db* mice. Moreover, GLUT4 protein expression significantly increased in liver of *db/db* mice treated with ML-SA8 by western blot (**Fig. 4c, d**) and immunofluorescence analysis (***Suppl. Fig.8d***), along with enhanced PM translocation (***Suppl. Fig.8e***). Notably, PM and post-PM fractions (PM-free supernatants) were isolated from skeletal muscle and adipose tissues, and quantification of the PM/Input ratio showed that ML-SA8 treatment increased GLUT4 PM translocation in skeletal muscle (***Suppl. Fig. 8f***) and adipose tissue (***Suppl. Fig. 8g***). Furthermore, administration of SA8 (4 and 8 mg/kg) significantly reduced fasting blood glucose (FBG) levels (**Fig. 4f**), HbA1c levels (**Fig. 4g**) and random blood glucose (RBG) levels (***Suppl. Fig.9a***), oral glucose tolerance tests (OGTT) (**Fig. 4h**) and insulin tolerance test (ITT) (**Fig. 4i**). These data suggest that ML-SA8 reduced plasma glucose levels and increased the insulin sensitivity in *db/db* mice. Histological staining analysis showed that liver tissues in DB+V mice exhibited hepatic steatosis, lipid droplet accumulation, hepatocyte swelling and ballooning (**Fig. 4j**). In contrast, administration of SA8 (4 mg/kg) in DB mice was sufficient to restore a normal, well-defined, typical hepatocyte morphology (**Fig. 4j**). Moreover, in the Periodic Acid-Schiff (PAS) staining used for glycogen deposition analysis, the glycogen levels in liver tissues of DB+SA8 group displayed a significant elevation compared with DB+V group (**Fig. 4j**), indicating that SA8 prevented reduced glycogen synthesis in *db/db* mice. Furthermore, ML-SA8 (4 mg/kg) treatment reduced serum alanine aminotransferase (ALT) (**Fig. 4k)** and aspartate aminotransferase (AST) levels (**Fig. 4l**) and increased plasma insulin concentration (***Suppl. Fig.9b***). Notably, ML-SA8 treatment had no effects on body weight (***Suppl. Fig.9c***), total cholesterol (TC) (***Suppl. Fig.9d***), serum triglyceride (TG) (***Suppl.Fig.9e***) and serum creatinine (***Suppl.Fig.9f***). Collectively, these findings indicate that ML-SA8 alleviates the hallmarks of diabetic phenotypes in *db/db* mice.

## Discussion

Lysosomes, a cellular hub in metabolism, protein degradation, and nutrient sensing, play an indispensable role in numerous metabolic pathways and maintaining cellular homeostasis [60]. Emerging evidence implicates lysosomal calcium signaling as a pivotal yet understudied modulator of metabolic homeostasis. In the current study, we found that genetic or pharmacological activation of the lysosomal Ca^2+^ channel-TRPML1 exhibited glucose-lowering effects in both *in vitro* and *in vivo* diabetic models. TRPML1 activation led to AMPK phosphorylation via a Ca^2+^-CaMKKβ-dependent mechanism and promoted GLUT4 translocation, thereby facilitating glucose influx to reduce hyperglycemia and other phenotypes of T2DM (**Fig. 4m**). Hence, our proof-of-concept study highlighted TRPML1 as a functional therapeutic target for T2DM and provided evidence that small-molecule TRPML1 agonists may have broad therapeutic potential.

Lysosomal Ca^2+^ is a key regulator for various lysosomal functions i.e., lysosomal exocytosis, membrane trafficking, catabolite export, ion homeostasis and nutrient sensing [61]. Ca^2+^ in the lysosomal lumen is about ∼ 0.5 mM, approximately 5,000-fold higher than cytosolic Ca^2+^ (∼100 nM). TRPML1 is predominantly localizes on the membranes of late endosomes and lysosomes (LELs) in all mammalian cell types [62]. The related TRPML2 and TRPML3 channels are also permeable to Ca^2+^, but are more restrictively expressed [62]. TRPML channels play essential roles in various lysosomal functions and respond to a variety of stimuli, such as pH, nutrients and cellular stress, as well as to small molecules [46], suggesting that TRPML1 activities can be specifically modulated. The synthetic agonists (ML-SAs) and inhibitors (ML-SIs) that specifically targeting TRPML1 directly bind to TRPML1 protein in the atomic-resolution co-structures [30, 31]. Notably, the concentration of ML-SAs used in this study has no cytotoxicity towards the cell lines we used including HepG2, HeLa and HAP1 cells (***Suppl. Fig.10a, b***). Given that *in vitro* experiments were performed primarily in transformed cell lines, and future studies in primary hepatocytes could provide further insights for these findings.

Although GLUT4 is expressed at a relatively low level in the liver compared with other tissues e.g. skeletal muscle, adipose, and heart, and is not the predominant hepatic glucose transporter (where GLUT2 predominates)[38], accumulating evidences indicates that GLUT4 plays a functional role in hepatic glucose homeostasis [40, 63–65]. For example, GLUT4 mediates glucose uptake in hepatic stellate cells [66]. Following ischemia-reperfusion injury, hepatic GLUT expression is dramatically upregulated, whereas GLUT2 is not. The increased GLUT4 localizes to the peripheral membranes of hepatocytes, where it enhances glucose uptake and promotes glycogen deposition [40, 67]. Furthermore, GLUT4-null mice exhibit compensatory changes in hepatic glucose and lipid metabolism, including increased hepatic glucose uptake and triglycerides conversion [68]. Collectively, these studies indicate a clear physiological role for GLUT4 in the liver. Under basal conditions, GLUT4 resides intracellularly and translocates to the PM upon insulin or other stimuli to drive glucose uptake[69]. Given that GLUT4 mediates AMPK-dependent glucose uptake via translocation [37], we examined the effect of ML-SA8 on GLUT4 localization and expression. While our findings support ML-SA-mediated regulation of GLUT4, the contribution of GLUT2 cannot be excluded. Moreover, ML-SA8 upregulated GLUT4 levels and PM translocation in skeletal muscle and adipose tissue of *db/db* mice (***Suppl.Fig.8b-f***), suggesting that the systemic glycemic recovery should be attributed to the multi-tissue effects of ML-SA8, rather than a liver-specific phenomenon. Regarding how GLUT4 expression is upregulated by TRPML1, Ca²⁺ from TRPML1 has been reported to increases its transcription. Two mechanisms have been proposed: the CaMKII pathway, where Ca²⁺ promotes histone acetylation and binding of myocyte enhancer factor 2 (MEF2) to the GLUT4 promoter [70] and the calcineurin pathway, where Ca²⁺-activated calcineurin dephosphorylates MEF2 or nuclear factor of activated T cells (NFAT) [71]. Given our TRPML1-CaMKKβ-AMPK findings, how CaMKII, calcineurin, CaMKKβ, or AMPK contributes to ML-SA8-TRPML1-driven transcription remains to be determined.

The MiT/TFE family includes TFEB, TFE3, MITF and TFEC. TFEB, TFE3 and MITF, but not TFEC are master regulators of lysosome and autophagy by controlling the “coordinated lysosomal expression and regulation” (CLEAR) network, covering genes associated to lysosomal exocytosis and biogenesis, and autophagy [55, 72, 73]. TFEB/TFE3/MITF are phosphorylated by MTOR (mechanistic target of rapamycin kinase) and then bind to YWHA/14-3-3 in the cytosol inactively [74, 75]. Whereas, under stress conditions i.e. starvation, TFEB/TFE3/MITF are dephosphorylated and actively translocate into nucleus, promoting the expression of genes [56, 76, 77]. TRPML1-released Ca^2+^ release reportedly activates calcineurin, which binds and dephosphorylates TFEB, thus promoting its nuclear translocation [77]. Interestingly, we found that AMPK is not required for TFEB nuclear translocation induced by ML-SA8 (***Suppl. Fig.7*d, e**). Furthermore, by using TFEB/TFE3/MITF TKO cells we found that knockout TFEB/TFE3/MITF has no effect on ML-SA8-induced AMPK activation (***Suppl. Fig.7*a-c**). These results suggest that TRPML1-mediated activation of TFEB and AMPK may be through distinct independent pathways.

In summary, we show a novel AMPK activation pathway via the lysosomal TRPML1-Ca²⁺-CaMKKβ axis. Pharmacological activation of TRPML1 can efficiently diminish diabetic phenotypes in both *in vitro* and *in vivo* models by targeting AMPK. Hence, targeting lysosomal Ca^2+^ channels i.e. TRPML1 may represent a promising approach to treat T2DM and related metabolic diseases.

## Methods and materials

### Cell lines and cell culture

HeLa (RRID: CVCL-0030), HepG2 (RRID: CVCL_A8FT) and C2C12 (RRID: CVCL-0188) cells were purchased from American Type Culture Collection (ATCC). Huh7 (RRID: RRID: CVCL-0336) cells were purchased from Thermo Scientific. HepG2, Huh7 and C2C12 cells were cultured in Dulbecco’s modified Eagle’s medium (DMEM; Thermo Scientific, 11195-065) supplemented with 10% fetal bovine serum (Thermo Scientific, 10091148) and 1% Penicillin-Streptomycin (PS; Solarbio, P1400). HeLa cells were maintained in modified Eagle’s medium (MEM; Thermo Scientific, 1964643) supplemented with 10% FBS and 1% PS. GFP-TFEB HeLa stable, HAP1 (RRID: CVCL-Y019, Horizon Company), *TRPML1* KO HAP1, HEK293-GCaMP7-TRPML1 and *ATG5* KO HAP1 cells were maintained in our laboratory as previously described [45, 56, 78, 79] and cultured in Iscove’s Modified Dulbecco’s Medium (IMDM; Gibco, 6123179) supplemented with 10% FBS and 1% PS. All cells were used after in-house authentication for mycoplasma contamination and maintained at 37 °C in a humidified 5% CO_2_ incubator. Cells with passage number less than 20 were used in this study.

### Construction of insulin resistance (IR) model

PA (Sigma-Aldrich, P0500) was applied to construct the IR model [80]. Briefly, 9.2 mg PA was completely dissolved in 1ml of 0.1 M NaOH by warming to 80 ℃ until becoming transparent and clear. The PA solution was further diluted in fatty acid-free bovine serum albumin solution (BSA, 0.325 g BSA dissolved in 8 ml of 0.9 % NaCl by heating at 45℃). The two solutions were then mixed with gentle agitation to get final PA stock solution (4 mM), which was further sterilized by passing through 0.22 μm filters, aliquoted and stored at -20 ℃. Cells were cultured at approximately 70% confluency and treated with PA (0.25 mM, 12 h) to construct IR model, followed by chemical treatment as required conditions.

### Plasmid transfection

Plasmids TRPML1-mCherry and GFP-TRPML1 were maintained in our laboratory [81]. Cells were seeded into 6-well plates and transfected with plasmids using Lipofectamine 3000 (Thermo Scientific, 2163785). The efficiency of the transfection was validated by Western blotting or Q-PCR.

### RNA interference

The siRNA sequences targeting human CaMKKβ (5ʹ-GGA CCA UCU GUA CAU GGU GUU CGA A-3ʹ) were chemically synthesized by GenePharma. HepG2 cells were seeded into 6-well plates and transfected with siRNA using Lipofectamine RNAi-Max reagent (Thermo Fisher Scientific, 13778150). The efficiency of the siRNA knockdown was assayed by Q-PCR.

### RNA extraction and Q-PCR

Total RNA was isolated from cells using Trizol (Thermo Scientific, 191002), and cDNA was generated using the GoScript Reverse Transcription System (Promega, 0000316057) according to the manufacturer’s protocol. Q-PCR was performed using the SYBR Green Mix (TOYOBO, 563700) in CFX Connect Optics (Bio-Rad). The changes in the mRNA expression of target genes were normalized to that of the housekeeping gene HPRT. Primer sequences used in this study are listed below:

*HPRT*; For 5ʹ-TGGCGTCGTGATTAGTGATG-3ʹ, Rev 5ʹ-CTGTTCTCGTCCAGCA GACACT-3ʹ

*CaMKKβ*; For 5ʹ- TCCAGACCACGACATAG -3ʹ, Rev 5ʹ-CAGGGGTGCAGCTTGATTTC-3ʹ

*TRPML1*; For 5ʹ- AGTGCCTGTTCTCGCTCATC; Rev 5ʹ-GGGATGCTTGATGGTGTCGT

### GCaMP7 Ca^2+^ imaging

GCaMP7 Ca^2+^ imaging was performed in HEK293 cells stably expressing GCaMP7-TRPML1, a lysosome-targeted genetically-encoded Ca^2+^ sensor [78]. Cells were excited at 488 nm, and the emission fluorescence was recorded at 510 nm using a FlexStation 3 scanning fluorometer. Fluorescence intensity at 488 nm excitation (F₄₈₈) was acquired at regular intervals under controlled temperature (e.g., 37 °C) and continuous perfusion with appropriate buffer. Data were normalized to baseline fluorescence (F₀) and presented as ΔF/F₀.

### Immunofluorescence and confocal imaging

For immunofluorescence detection of GLUT4, cells were grown and stained on coverslips in 24-well plates. 4% paraformaldehyde (Solarbio, P1110) was added to each well and cells were fixed for 15 min at room temperature. 0.3% Triton X-100 (Sigma-Aldrich, T8787) was used for permeabilization for 10 min, followed by 1 h blocking with 1% bovine serum albumin (Sigma-Aldrich, WXBC2612V) in phosphate-buffered saline (PBS; Thermo Scientific, 21600069). The cells were incubated overnight at 4°C with anti-GLUT4 antibodies (Abcam, AB33780) (1:1000). After washing for 5 times with PBS, secondary antibodies conjugated to Alexa Fluor -488 (Thermo Scientific, 1705869) (1:1000) were added and incubated for 2 h in a dark place. DAPI (Sigma-Aldrich, D9542) was added to stain nuclei for 15 min and briefly washed for 3 times. Coverslips were then mounted with Fluoromount-G (Southern Biotech) and images were acquired with an Olympus or Zeiss confocal microscope. The fluorescence intensity of GLUT4 membrane staining was quantified by performing a line scan across one or two cells and measuring fluorescence along the line.

### Measurement of cellular glucose uptake

Cellular glucose uptake was measured using a fluorescence-labeled glucose analogue-2-NBDG (MCE, 186689-07-6) [44]. Briefly, cells were seeded about 70% confluency and treated with chemicals as required conditions. Cells were then washed and fresh DMEM (no glucose, no FBS, no phenol red, Solarbio, D6540) containing 50 μM 2-NBDG was added for 30 min, followed by PBS wash three times. The fluorescence was visualized with an Olympus IX-73 inverted microscope. Fluorescence intensity was quantified using Image J software.

### Measurement of extracellular glucose consumption

Glucose consumption in the medium was measured using a glucose consumption kit (Nanjing Jiancheng Bioengineering Institute, A154-1-1) [82]. Cells were seeded in a 6-well plate at a density of 1 × 10^6^ cells/ml and treated with chemicals as required conditions in DMEM (no FBS, no phenol red) and supplemented with 10 mM glucose. Following the manufacturer’s protocol, 2.5 μL of sample (medium/standard) was mixed with 250 μL of working solution a 96-well plate. The mixture was then incubated at 37 °C for 10 min and the absorbance was measured at 505 nm using a spectrophotometer VARIOSKAN LUX (Thermo Scientific). Glucose concentration was calculated as: [Medium Glucose] = [(Sample_absorbance_-blank_absorbance_) / (Standard_absorbance_-blank_absorbance_)] × 5.55 mmol/L × X, where 5.55 mmol/L is the concentration of the glucose standard, and X is the dilution factor.

### Measurement of ATP/ADP ratio

The ATP/ADP ratio of cells was measured with an ATP/ADP Ratio Chemiluminescence Assay Kit (Elabscience, E-BC-F004) following the manufacturer’s instructions. Briefly, approximately 1 × 10^6^ cells were incubated with nucleotide-releasing buffer (150 μL) for 10 min in boiling water. Then, 100 μL of ATP monitoring enzyme was added into a 96-well plate to measure the background luminescence (L1). After adding 20 μL cell lysate, luminescence (L2) was recorded immediately, with ATP=L2-L1. After 20 min, luminescence (L3) was determined. Following addition of 50 μL ADP convertase and incubation for 6 min at room temperature, luminescence (L4) was recorded, with ADP= L4-L3. The ATP/ADP ratio was calculated as (L2-L1)/(L4-L3).

### Protein extraction and Western blotting

Cells or tissues were lysed in RIPA buffer (Solarbio, R0010) supplemented with 1x protease inhibitors cocktail (Sigma-Aldrich, P8340) (1: 100) and 1 x phosphatase inhibitor cocktail (Abcam, GR304037-28) (1: 100) on ice for 20 min. The membrane protein fraction was extracted using a membrane and cytoplasm protein extraction kit (Beyotime, P0033, Shanghai, China). A total of 2 × 10⁷ cells or approximately 30 mg of tissues were processed following the manufacturer’s protocol. Protein samples were extracted and separated by sodium dodecyl sulfate-polyacrylamide gel electrophoresis gels, followed by transferring to polyvinylidene difluoride membranes (Merck, R7DA8778E). The membranes were blocked with 5% nonfat dry milk for 1 h and incubated with antibodies against AMPK (Cell Signaling Technology, 2532) (1:1000), p-AMPK (Cell Signaling Technology, 2535) (1:1000), ACC(Affinity Biosciences, AF6421)(1:1000), p-ACC (Affinity Biosciences, AF3421)(1:1000), GLUT4 (Abcam, AB33780) (1:1000), Vinculin (Abcam, AB8226) (1:2000), TFEB (Cell Signaling Technology, 4240S) (1:1000), TFE3 (HPA02300881) (1:1000), MITF (Abcam, AB20663) (1:1000) and GAPDH (Sigma-Aldrich, G9545) (1:10,000), respectively. Bound antibodies were detected horseradish peroxidase-conjugated anti-rabbit (Abcam, ab6721) or anti-mouse secondary antibodies (Abcam, ab6789) (1:10,000) in 5 % nonfat dry milk for 2 h at room temperature with agitation. Protein bands were visualized using enhanced chemiluminescence reagents (Thermo Scientific, 203-17071) in a Li-COR Biosciences Odyssey Fc system. Band intensities were quantified using the Image J software.

### Animals

Specific pathogen free (SPF) grade male C57BLKS/J mice (8-week-old) and male C57BLKS/J-Lepr KO (*db/db*) mice were purchased from GemPharmatech Limited Company (Nanjing, China). Throughout the manuscript, “DB” denotes genetically diabetic db/db mice. Animals were handled according to approved institutional animal care and use committee (IACUC) protocols (20241226208) of Zhejiang University of Technology. Mice were acclimated for one week prior to the experiment. After the acclimation period, mice were randomly assigned to experimental groups. At the end of the experiments, all mice were euthanized by CO₂ inhalation followed by cervical dislocation to confirm death. Blood and liver tissues were rapidly collected for subsequent biochemical and molecular analyses. All experimental procedures were conducted in accordance with the Guide for the Care and Use of Laboratory Animals in the Zhejiang University of Technology, Hangzhou, China, and conformed to the National Institutes of Health Guide for Care and Use of Laboratory Animals.

### Histopathological analysis (H&E and PAS staining)

Liver tissues were fixed in 4 % paraformaldehyde buffer and embedded in paraffin. Liver slices were then deparaffinized, rehydrated, and stained with H&E (Solarbio, G1120) or PAS (Solarbio, G1281). All staining procedures were performed according to standard protocols followed by microscopy examination. The section images were obtained using the Olympus/Zeiss Fluorescence inverted microscope.

### Blood glucose and insulin measurements

Oral Glucose Tolerance Test (OGTT) and Insulin Tolerance Test (ITT) were performed on each group of mice. For OGTT, mice fasted for 10 h received oral glucose (1g/kg). For ITT, mice fasted for 4 h received i.p. injection of insulin (0.75 U /kg). Tail blood glucose was measured at 0, 30, 60, 90, and 120 min post-administration. Random blood glucose (RBG) was measured every four days (9:00–11:00 AM) to assess non-fasted status, minimizing circadian variation. Fasting blood glucose (FBG) was measured at baseline (Week 0) and endpoint (Week 6) after 6 h fast. Blood glucose was determined using a Sinocare glucometer, and plasma insulin by an ultrasensitive insulin ELISA kit (Solarbio, SEKM-0141). Area under the curve (AUC) was calculated for glucose response. Blood samples were collected through mice tail. After sterilizing the tail with an alcohol swab, a small incision was made at the tail vein tip after discarding the first droplet.

### Reagents

Chemicals used in this study including BAPTA-AM (Thermo Scientific, 1824047), EGTA-AM (Sigma-Aldrich, E3478), DFO (MCE, HY-B0988), TPEN (MCE, HY-100202), GPN (Santa Cruz Biotech, sc-252858), riluzole (Sigma-Aldrich, R116), PA (Sigma-Aldrich, P0500), insulin (Beyotime, P3376), metformin (Beyotime, S1741), CC (MCE, HY-13418A), STO609 (MCE, HY-19805), TG assay (Solarbio, BC0625), TC assay (Solarbio, BC1985), ALT assay (Solarbio, BC1555), AST assay (Solarbio, BC1565), HbA1c kit (Solarbio, BC5615), Creatinine assay (Nanjing Jiancheng Company, C011-2-1). ML-SA5, ML-SA8 and ML-SI5 were custom synthesized (available upon MTA request).

### Statistical analysis

Data are presented as mean ± s.e.m. from at least 3 independent experiments. Statistical comparisons were performed using a two tailed *Student’s t-test* (paired or unpaired as appropriate) or one-way/two-way ANOVA followed by Tukey’s or Dunnett’s post hoc test for multiple comparisons. A *P* value < 0.05 was considered statistically significant.

## Acknowledgements

This work was supported by a grant from National Natural Science Foundation of China (31600823 to D. L.). Additional support was provided by Calygene Biotechnology Inc. (XT [2016]008@), Lysoway Therapeutics, Inc. (KYY-HX-20210129) and Lyso-X Therapeutics (KYY-HX-20240478). The funders had no role in study design, data collection and analysis, decision to publish, or preparation of the manuscript. We appreciate the encouragement and assistance provided by Dr. Haoxing Xu and his laboratory members. The authors thanks Experimental Animal Center of Zhejiang University of Technology for providing the animal facilities and technical support for this study.

## Author contributions

J.Z., Data curation, Investigation, Methodology and Writing original draft; Z.P., Data curation, Validation, Investigation, Methodology and Writing original draft, review and editing; S.W., Data curation, Investigation, Methodology and Writing review and editing; H.Z., Data curation, Methodology, Writing review and editing; Q.W., Z.J, Y.W., Z.D., Investigation, Methodology; D.L., Conceptualization, Supervision, Funding acquisition, Validation, Investigation, Methodology, Project administration, Writing original draft, review and finalization.

## Data Availability Statement

Data available on request from the authors.

## Abbreviations

ALT: alanine aminotransferase
ACC: acetyl-CoA carboxylase
AMP: adenosine monophosphate
AMPK: AMP-activated protein kinase
ANOVA: analyses of variance
AST: aspartate aminotransferase
BAPTA-AM: 1,2-bis(2-aminophenoxy)ethane-N,N,N’,N’-tetraacetic acid tetrakis-acetoxymethyl ester
CaMKKβ: calcium/calmodulin-dependent protein kinase kinase β
DFO: deferoxamine mesylate
EGTA-AM: EGTA Acetoxymethyl ester
ER: endoplasmic reticulum
FBG: fasting blood glucose
FBS: fetal bovine serum
GLUT4: Glucose Transporter 4
GPN: Glycyl-L-phenylalanine 2-naphthylamide
HbA1c: hemoglobin A1c
H&E: hematoxylin-eosin
ITT: insulin tolerance test
IR: insulin resistance
KO: knockout
LKB1: liver kinase B1
MCOLN1/TRPML1: mucolipin 1
MET: metformin
ML-SA8: mucolipin-specific synthetic agonist 8
ML-SI5: mucolipin-specific synthetic inhibitor 5
OGTT: oral glucose tolerance test
PA: palmitic acid
PAS: periodic acid-schiff
PBS: phosphate-buffered saline
Q-PCR: quantitative real time polymerase chain reaction
RBG: random blood glucose
TFEB: transcription factor EB
TG: triglyceride
TC: total cholesterol
TPCs: two-pore channels
TPEN: N,N,N’,N’-tetrakis (2-pyridylmethyl) ethylene-diamine
T2DM: type 2 diabetes mellitus
ULK1: Unc like autophagy activating kinase 1
WT: wild-type
2-NBDG: 2-deoxy-2-[(7-nitro-2,1,3-benzoxadiazol-4-yl)amino]-d-glucose;

## Supplementary Figure Legends

**Suppl. Fig. 1. ML-SA8 activates AMPK in various cell lines**. **(a)** ML-SA8 (0.0003-3 μM) induced lysosomal Ca^2+^ release, measured with the GCaMP7 signal (F_470_), in GCaMP7-TRPML1-expressing HEK293 cells. (**b**) ML-SA8 dose-dependently activated lysosomal TRPML1 currents in GCaMP7-TRPML1-expressing HEK293 cells. (**c**) ML-SA8 (0.3-3 μM, 2 h) activated AMPK in HeLa cells in a dose-dependent manner analyzed by Western blotting. (**d**) Quantification of results shown in **a**. (n= 3). (**e**) ML-SA8 (0.3-3 μM, 2 h) induced AMPK activation in HAP1 cells. (**f**) Quantification of ratio of p-AMPK/AMPK protein levels shown in **c**. (n= 3 independent experiments). For all the panels, data are presented as mean ± s.e.m.; * *p* < 0.05, ** *p* < 0.01, *** *p* < 0.001, ANOVA.

**Suppl. Fig. 2. Multiple TRPML1 specific agonists activate AMPK in various cell lines**. Treatment of ML-SA5 (1 μM) and ML-SA8 (1 μM) for 2 h induced AMPK phosphorylation in (**a**) HepG2 **(b**) HeLa and (**c**) HAP1 cells analyzed by Western blotting. (**d**) Treatment with ML-SA8 (1 μM, 2 h) increased the p-ACC/total ACC ratio in HepG2, HeLa, and HAP1 cells. (**e**) ML-SA8 (1μM, 4 h) has no effect on the ADP/ATP ratio in HepG2 cells. Data are presented as mean ± s.e.m.; “ns” represents no significant difference.

**Suppl. Fig. 3. The effect of TPC2 agonist on AMPK activation**. (**a**) Riluzole (a TPC2 synthetic agonist, 200 μM) did not induce AMPK phosphorylation compared to ML-SA8 treatment in HepG2 cells. (**b**) The effect of Riluzole (200-800 μM, 2 h) on AMPK activation in HepG2 cells.

**Suppl. Fig. 4. The effect of ML-SA8 on GLUT4 mRNA and protein expression levels.** (**a**) QPCR analyzes the effect of ML-SA8 (1 μM, 4-24 h) on GLUT4 mRNA levels. (**b**) Western blotting analyzes the effect of ML-SA8 (1 μM, 24 h) on GLUT4 protein expression in IR-HepG2 cells. Data are presented as mean ± s.e.m.; \*\*\**p* < 0.001, ANOVA.

**Suppl. Fig. 5. ML-SA8 induced glucose influx in a dose-dependent manner in PA-induced insulin resistance (IR) models.** (**a**) Validation of IR-HepG2 model. HepG2 cells were treated with PA (0.25 mM,12 h) in complete medium, followed by insulin (INS, 100 nM, 0.5 h) treatment in glucose-, FBS-, and phenol red-free DMEM. Cells were then incubated with 2-NBDG (50 µM, 0.5 h). Scale bar, 40 μm. (**b**) Average ratio of the fluorescent intensity of 2-NBDG shown in **a**. from n= 3 independent experiments. (**c**)Western blotting analysis of AMPK phosphorylation status in IR-HepG2 cells induced by ML-SA8 (1 μM) or MET (2 mM) for 2 h. Vinculin as a loading control. (**d**) ML-SA8 (1 μM, 2-6 h) induced cellular glucose influx in IR-HepG2 cells in a time-dependent manner measured by 2-NBDG intensities. Scale bar, 40 μm. (**e**) Quantification of the fluorescent intensity of 2-NBDG shown in **d**. (n= 3). (**f**) ML-SA8 promotes glucose influx in various IR cell models. Huh7 or C2C12 cells were treated with PA (0.25 mM, 12 h) to construct IR-models, followed by the treatment of ML-SA8 (1 μM, 4 h), then incubated with 2-NBDG (50 µM, 0.5 h). (**g**) Quantitative analysis of the fluorescent intensity of 2-NBDG in **f**. (n= 3). For all the panels, data are presented as mean ± s.e.m.; ** *p* < 0.01, \*\*\**p* < 0.001, ANOVA.

**Suppl. Fig. 6. Chelation of Fe^2+^ or Zn^2+^ has no effect on ML-SA8-induced AMPK activation.** (**a**) Validation of *TRPML1* KO HAP1 cells by QPCR (**b**) BAPTA-AM inhibited ML-SA8-induced ACC phosphorylation in HepG2 cells. Cells were pretreated with BAPTA-AM (fast Ca^2+^ chelator, 20 μM) or EGTA-AM (slow Ca^2+^ chelator, 20 μM) for 1 h, then cotreated with ML-SA8 (1 μM, 2 h), p-ACC/total ACC was then evaluated by Western blotting. (**c**) BAPTA-AM (20 μM) and EGTA-AM (20 μM) inhibited Ca²⁺ induced by ionomycin [83], (5 μM, which directly mediates extracellular Ca²⁺ influx) and thapsigargin [84] (2 μM, which indirectly elevates intracellular Ca²⁺ by inhibiting SERCA to deplete endoplasmic reticulum calcium stores). (**d**) Effects of DFO (a Fe^2+^ chelator) and TPEN (a Zn^2+^ chelator) on ML-SA8-induced AMPK activation. Western blot analysis of p-AMPK/total AMPK in HepG2 cells cotreated with ML-SA8 (1 μM) and DFO (10 μM), or TPEN (10 μM) for 2 h. (**e**) Ratio of p-AMPK vs. total AMPK as shown in **d**. from n= 4 independent experiments. (**f**) The effects of DFO and TPEN on ML-SA8-induced cellular glucose uptake. IR-HepG2 cells treated with ML-SA8 (1 μM, 4 h) with or without DFO (10 μM) or TPEN (10 μM) for 4 h. Scale bar, 40 μm. (**g**) Quantification of 2-NBDG in **f**. from n= 3 independent experiments. For all the panels, data are presented as mean ± s.e.m.; “ns” represents no significant difference, ANOVA.

**Suppl. Fig. 7. TFEB-independence of ML-SA8-induced AMPK activation.** (**a**) Western blot analysis of *TFEB/TFE3/MITF* triple KO (TKO) efficiency in HeLa cells. (**b**) ML-SA8-induced AMPK activation in *TFEB/TFE3/MITF* TKO HeLa cells. WT and TKO HeLa cells were treated with ML-SA8 (1 μM) for 2 h. (**c**) Quantitative analysis of p-AMPK/total AMPK as shown in **b**. (n= 3). (**d**) Differential effects of ML-SI, CC and STO609 on ML-SA8-induced TFEB nuclear translocation. HeLa GFP-TFEB stable cells was pretreated with ML-SI5 (3 μM), CC (10 μM), or STO609 (10 μM) for 1 h, followed by cotreatment with ML-SA8 (1 μM, 4 h). Nuclei were counterstained with DAPI (blue). Scale bar, 10 μm. (**e**) Quantification of results shown in **d**. (N=30 randomly-selected cells from at least 3 independent experiments). For all the panels, data are presented as mean ± s.e.m.; ** *p* < 0.01, \*\*\**p* < 0.001. “ns” represents no significant difference.

**Suppl. Fig. 8. The effects of ML-SA8 on AMPK activation and GLUT4 in *db/db* mice**. *db/db* mice were i.p. injected with ML-SA8 (4 mg/kg) or vehicle (5% DMSO, 90% PEG300, 5% ddH_2_O) for 6 weeks. **(a)** liver **(b)** adipose and **(c)** skeletal muscle tissues were collected and subjected to detect p-AMPK/total AMPK by Western blotting. GLUT4 expression was examined in liver tissues of *db/db* mice with 6-weeks injection of ML-SA8 (4mg/kg) by immunofluorescence. Scale bar 40 μm. GLUT4 PM translocation was examined in (**e**) liver (**f**) skeletal muscle (**g**) adipose tissues of *db/db* mice with or without 6-weeks injection of ML-SA8 (4mg/kg). Vinculin served as the loading control for the Input and Post-PM fractions. Ponceau S (Po-S) staining was used as the loading control for the PM fractions.

**Suppl. Fig. 9. The effects of ML-SA8 on phenotypes of *db/db* mice**. *db/db* mice were i.p. injected with ML-SA8 (4 mg/kg) or vehicle (5% DMSO, 90% PEG300, 5% ddH_2_O) for 6 weeks. (**a**) Blood glucose was detected by glucometer every four days. AUC values were calculated. (**b**) Daily mice body weight (**c**) serum insulin (**d**) TC (**e**) TG and (**f**) Creatinine in various groups. In all panels, data are presented as mean ± s.e.m. \**p* < 0.05, \*\**p* < 0.01, *** *p*< 0.001, ANOVA.

**Suppl. Fig. 10. ML-SA8 has no cytotoxic effect on various cell lines**. The cytotoxic effect of ML-SA8 (0-30 μM, 4 h) in (**a**) HepG2, HeLa and HAP1 cells and (**b**) IR-HepG2 and IR-HAP1 cells. In all panels, data are presented as mean ± s.e.m. \**p* < 0.05, \*\**p* < 0.01.

## References

1. Ahmad, E., et al., Type 2 diabetes. Lancet, 2022. 400(10365): p. 1803–1820.

2. Sun, H., et al., IDF Diabetes Atlas: Global, regional and country-level diabetes prevalence estimates for 2021 and projections for 2045. Diabetes Res Clin Pract, 2022. 183: p. 109119.

3. González, A., et al., AMPK and TOR: The Yin and Yang of Cellular Nutrient Sensing and Growth Control. Cell Metab, 2020. 31(3): p. 472–492.

4. Hardie, D.G., Keeping the home fires burning: AMP-activated protein kinase. J R Soc Interface, 2018. 15(138).

5. Woods, A., et al., Ca2+/calmodulin-dependent protein kinase kinase-beta acts upstream of AMP-activated protein kinase in mammalian cells. Cell Metab, 2005. 2(1): p. 21–33.

6. Ross, F.A., T.E. Jensen, and D.G. Hardie, Differential regulation by AMP and ADP of AMPK complexes containing different gamma subunit isoforms. Biochem J, 2016. 473(2): p. 189–99.

7. Hawley, S.A., et al., Complexes between the LKB1 tumor suppressor, STRAD alpha/beta and MO25 alpha/beta are upstream kinases in the AMP-activated protein kinase cascade. J Biol, 2003. 2(4): p. 28.

8. Clapham, D.E., Calcium signaling. Cell, 2007. 131(6): p. 1047–58.

9. Scotto Rosato, A., et al., TRPML1 links lysosomal calcium to autophagosome biogenesis through the activation of the CaMKKβ/VPS34 pathway. Nat Commun, 2019. 10(1): p. 5630.

10. Zhong, X.Z., et al., BK channel agonist represents a potential therapeutic approach for lysosomal storage diseases. Sci Rep, 2016. 6: p. 33684.

11. Somogyi, A., et al., The synthetic TRPML1 agonist ML-SA1 rescues Alzheimer-related alterations of the endosomal-autophagic-lysosomal system. J Cell Sci, 2023. 136(6): p. JCS259875.

12. Bae, M., et al., Activation of TRPML1 clears intraneuronal Abeta in preclinical models of HIV infection. J Neurosci, 2014. 34(34): p. 11485–503.

13. Bargal, R., et al., Identification of the gene causing mucolipidosis type IV. Nat Genet, 2000. 26(1): p. 118–23.

14. Pu, J., Targeting the lysosome: Mechanisms and treatments for nonalcoholic fatty liver disease. J Cell Biochem, 2022. 123(10): p. 1624–1633.

15. Arden, C., et al., Autophagy and lysosomal dysfunction in diabetes and its complications. Trends Endocrinol Metab, 2024. 35(12): p. 1078–1090.

16. Spampanato, C., et al., Transcription factor EB (TFEB) is a new therapeutic target for Pompe disease. EMBO Mol Med, 2013. 5(5): p. 691–706.

17. Feng, X., et al., Activation of lysosomal Ca2+ channels mitigates mitochondrial damage and oxidative stress. J Cell Biol, 2025. 224(1).

18. Yu, L., et al., Small-molecule activation of lysosomal TRP channels ameliorates Duchenne muscular dystrophy in mouse models. Sci Adv, 2020. 6(6): p. eaaz2736.

19. Chen, C.C., et al., A small molecule restores function to TRPML1 mutant isoforms responsible for mucolipidosis type IV. Nat Commun, 2014. 5: p. 4681.

20. Shi, W., et al., Saikosaponin-d inhibits proliferation by up-regulating autophagy via the CaMKKβ– AMPK–mTOR pathway in ADPKD cells. Molecular and Cellular Biochemistry, 2018. 449(1-2): p. 219–226.

21. Vahidi Ferdowsi, P., et al., TRPV1 Activation by Capsaicin Mediates Glucose Oxidation and ATP Production Independent of Insulin Signalling in Mouse Skeletal Muscle Cells. Cells, 2021. 10(6): p. 1560.

22. Morgan, A.J., et al., Molecular mechanisms of endolysosomal Ca2+ signalling in health and disease. Biochem J, 2011. 439(3): p. 349–74.

23. Prins, D. and M. Michalak, Organellar calcium buffers. Cold Spring Harb Perspect Biol, 2011. 3(3): p. a004069.

24. Hawley, S.A., et al., Characterization of the AMP-activated protein kinase kinase from rat liver and identification of threonine 172 as the major site at which it phosphorylates AMP-activated protein kinase. J Biol Chem, 1996. 271(44): p. 27879–87.

25. Hu, M., et al., The ion channels of endomembranes. Physiol Rev, 2024. 104(3): p. 1335–1385.

26. Wang, X., et al., TPC proteins are phosphoinositide-activated sodium-selective ion channels in endosomes and lysosomes. Cell, 2012. 151(2): p. 372–83.

27. Du, W., et al., Lysosomal Zn(2+) release triggers rapid, mitochondria-mediated, non-apoptotic cell death in metastatic melanoma. Cell Rep, 2021. 37(3): p. 109848.

28. Zhou, G., et al., Role of AMP-activated protein kinase in mechanism of metformin action. J Clin Invest, 2001. 108(8): p. 1167–74.

29. Sahoo, N., et al., Gastric Acid Secretion from Parietal Cells Is Mediated by a Ca2+ Efflux Channel in the Tubulovesicle. Developmental Cell, 2017. 41(3): p. 262–273.e6.

30. Schmiege, P., et al., Human TRPML1 channel structures in open and closed conformations. Nature, 2017. 550(7676): p. 366–370.

31. Schmiege, P., M. Fine, and X. Li, Atomic insights into ML-SI3 mediated human TRPML1 inhibition. Structure, 2021. 29(11): p. 1295–1302 e3.

32. Zhang, X., et al., Agonist-specific voltage-dependent gating of lysosomal two-pore Na(+) channels. Elife, 2019. 8.

33. Beltran-Parrazal, L. and A. Charles, Riluzole inhibits spontaneous Ca2+ signaling in neuroendocrine cells by activation of K+ channels and inhibition of Na+ channels. Br J Pharmacol, 2003. 140(5): p. 881–8.

34. Albertini, C., et al., Riluzole–Rasagiline Hybrids: Toward the Development of Multi-Target-Directed Ligands for Amyotrophic Lateral Sclerosis. ACS Chemical Neuroscience, 2022. 13(15): p. 2252–2260.

35. Dolfi, S.C., et al., Riluzole exerts distinct antitumor effects from a metabotropic glutamate receptor 1-specific inhibitor on breast cancer cells. Oncotarget, 2017. 8(27): p. 44639–44653.

36. Kakoti, B.B., et al., AMPK pathway: an emerging target to control diabetes mellitus and its related complications. J Diabetes Metab Disord, 2024. 23(1): p. 441–459.

37. Addinsall, A.B., et al., Electrical stimulated GLUT4 signalling attenuates critical illness-associated muscle wasting. J Cachexia Sarcopenia Muscle, 2022. 13(4): p. 2162–2174.

38. Karim, S., D.H. Adams, and P.F. Lalor, Hepatic expression and cellular distribution of the glucose transporter family. World J Gastroenterol, 2012. 18(46): p. 6771–81.

39. He, C., et al., Natural exosomes-like nanoparticles in mung bean sprouts possesses anti-diabetic effects via activation of PI3K/Akt/GLUT4/GSK-3β signaling pathway. J Nanobiotechnology, 2023. 21(1): p. 349.

40. Kurabayashi, A., et al., Murine remote ischemic preconditioning upregulates preferentially hepatic glucose transporter-4 via its plasma membrane translocation, leading to accumulating glycogen in the liver. Life Sci, 2022. 290: p. 120261.

41. Hu, W., et al., Hirsutine ameliorates hepatic and cardiac insulin resistance in high-fat diet-induced diabetic mice and in vitro models. Pharmacol Res, 2022. 177: p. 105917.

42. Samuel, V.T. and G.I. Shulman, Mechanisms for insulin resistance: common threads and missing links. Cell, 2012. 148(5): p. 852–71.

43. Xiao, H., et al., Gentiopicroside targets PAQR3 to activate the PI3K/AKT signaling pathway and ameliorate disordered glucose and lipid metabolism. Acta Pharm Sin B, 2022. 12(6): p. 2887–2904.

44. Yamada, K., et al., Measurement of glucose uptake and intracellular calcium concentration in single, living pancreatic beta-cells. J Biol Chem, 2000. 275(29): p. 22278–83.

45. Qi, J., et al., MCOLN1/TRPML1 finely controls oncogenic autophagy in cancer by mediating zinc influx. Autophagy, 2021. 17(12): p. 4401–4422.

46. Dong, X.-P., et al., The type IV mucolipidosis-associated protein TRPML1 is an endolysosomal iron release channel. Nature, 2008. 455(7215): p. 992–996.

47. Marcelo, K.L., A.R. Means, and B. York, The Ca 2+ /Calmodulin/CaMKK2 Axis: Nature’s Metabolic CaMshaft. Trends in Endocrinology & Metabolism, 2016. 27(10): p. 706–718.

48. Medina, D.L., et al., Lysosomal calcium signalling regulates autophagy through calcineurin and TFEB. Nat Cell Biol, 2015. 17(3): p. 288–99.

49. Hay, J.C., Calcium: a fundamental regulator of intracellular membrane fusion? EMBO reports, 2007. 8(3): p. 236–240.

50. Radford, R.J. and S.J. Lippard, Chelators for investigating zinc metalloneurochemistry. Curr Opin Chem Biol, 2013. 17(2): p. 129–36.

51. Pramanik, S., et al., Cell Permeable Imidazole-Desferrioxamine Conjugates: Synthesis and In Vitro Evaluation. Bioconjug Chem, 2019. 30(3): p. 841–852.

52. Hawley, S.A., et al., Calmodulin-dependent protein kinase kinase-beta is an alternative upstream kinase for AMP-activated protein kinase. Cell Metab, 2005. 2(1): p. 9–19.

53. Kim, J., et al., AMPK activators: mechanisms of action and physiological activities. Exp Mol Med, 2016. 48(4): p. e224.

54. Tokumitsu, H., et al., STO-609, a specific inhibitor of the Ca(2+)/calmodulin-dependent protein kinase kinase. J Biol Chem, 2002. 277(18): p. 15813–8.

55. Napolitano, G. and A. Ballabio, TFEB at a glance. J Cell Sci, 2016. 129(13): p. 2475–81.

56. Zhang, X., et al., MCOLN1 is a ROS sensor in lysosomes that regulates autophagy. Nat Commun, 2016. 7: p. 12109.

57. Paquette, M., et al., AMPK-dependent phosphorylation is required for transcriptional activation of TFEB and TFE3. Autophagy, 2021. 17(12): p. 3957–3975.

58. Sardiello, M., et al., A gene network regulating lysosomal biogenesis and function. Science, 2009. 325(5939): p. 473–7.

59. Suriano, F., et al., Novel insights into the genetically obese (ob/ob) and diabetic (db/db) mice: two sides of the same coin. Microbiome, 2021. 9(1): p. 147.

60. Lawrence, R.E. and R. Zoncu, The lysosome as a cellular centre for signalling, metabolism and quality control. Nat Cell Biol, 2019. 21(2): p. 133–142.

61. Xu, H. and D. Ren, Lysosomal physiology. Annu Rev Physiol, 2015. 77: p. 57–80.

62. Cheng, X., et al., Mucolipins: Intracellular TRPML1-3 channels. FEBS Lett, 2010. 584(10): p. 2013–21.

63. Rossetti, L., et al., Peripheral but not hepatic insulin resistance in mice with one disrupted allele of the glucose transporter type 4 (GLUT4) gene. J Clin Invest, 1997. 100(7): p. 1831–9.

64. Fan, X., et al., Ameliorative effects of mangiferin derivative TPX on insulin resistance via PI3K/AKT and AMPK signaling pathways in human HepG2 and HL-7702 hepatocytes. Phytomedicine, 2023. 114: p. 154740.

65. Jiang, Y., et al., The role of STAT3/VAV3 in glucolipid metabolism during the development of HFD-induced MAFLD. Int J Biol Sci, 2024. 20(6): p. 2027–2043.

66. Tang, Y. and A. Chen, Curcumin prevents leptin raising glucose levels in hepatic stellate cells by blocking translocation of glucose transporter-4 and increasing glucokinase. Br J Pharmacol, 2010. 161(5): p. 1137–49.

67. Kim, S., et al., Exendin-4 Improves Nonalcoholic Fatty Liver Disease by Regulating Glucose Transporter 4 Expression in ob/ob Mice. Korean J Physiol Pharmacol, 2014. 18(4): p. 333–9.

68. Ranalletta, M., et al., Altered hepatic and muscle substrate utilization provoked by GLUT4 ablation. Diabetes, 2005. 54(4): p. 935–43.

69. Huang, S. and M.P. Czech, The GLUT4 glucose transporter. Cell Metab, 2007. 5(4): p. 237–52.

70. Ojuka, E.O., V. Goyaram, and J.A. Smith, The role of CaMKII in regulating GLUT4 expression in skeletal muscle. Am J Physiol Endocrinol Metab, 2012. 303(3): p. E322–31.

71. Ryder, J.W., et al., Skeletal muscle reprogramming by activation of calcineurin improves insulin action on metabolic pathways. J Biol Chem, 2003. 278(45): p. 44298–304.

72. Martini-Stoica, H., et al., The Autophagy-Lysosomal Pathway in Neurodegeneration: A TFEB Perspective. Trends Neurosci, 2016. 39(4): p. 221–34.

73. Sardiello, M. and A. Ballabio, Lysosomal enhancement: a CLEAR answer to cellular degradative needs. Cell Cycle, 2009. 8(24): p. 4021–2.

74. Martina, J.A. and P. Rosa, Protein phosphatase 2A stimulates activation of TFEB and TFE3 transcription factors in response to oxidative stress. Journal of Biological Chemistry, 2018. 293: p. jbc.RA118.003471-.

75. Napolitano, G., et al., mTOR-dependent phosphorylation controls TFEB nuclear export. Nat Commun, 2018. 9(1): p. 3312.

76. Puertollano, R., et al., The complex relationship between TFEB transcription factor phosphorylation and subcellular localization. EMBO J, 2018. 37(11).

77. Medina, et al., Lysosomal calcium signalling regulates autophagy through calcineurin and TFEB. 2015.

78. Li, D., et al., Sulforaphane Activates a lysosome-dependent transcriptional program to mitigate oxidative stress. Autophagy, 2021. 17(4): p. 872–887.

79. Shen, D., et al., Lipid storage disorders block lysosomal trafficking by inhibiting a TRP channel and lysosomal calcium release. Nat Commun, 2012. 3: p. 731.

80. Lee, J.Y., H.K. Cho, and Y.H. Kwon, Palmitate induces insulin resistance without significant intracellular triglyceride accumulation in HepG2 cells. Metabolism, 2010. 59(7): p. 927–34.

81. Samie, M., et al., A TRP channel in the lysosome regulates large particle phagocytosis via focal exocytosis. Dev Cell, 2013. 26(5): p. 511–24.

82. Kabasakalian P Fau - Kalliney, S., A. Kalliney S Fau - Westcott, and A. Westcott, Enzymatic blood glucose determination by colorimetry of N,N-diethylaniline-4 aminoantipyrine. 20(5): p. 606–607.

83. Phillippe, M., et al., Ionomycin-stimulated phasic myometrial contractions. Am J Physiol, 1995. 269(4 Pt 1): p. E779–85.

84. Treiman, M., C. Caspersen, and S.B. Christensen, A tool coming of age: thapsigargin as an inhibitor of sarco-endoplasmic reticulum Ca(2+)-ATPases. Trends Pharmacol Sci, 1998. 19(4): p. 131–5.

