## Supplementary figures and images for "Targeting Lysosomal MCOLN1/TRPML1 Ion Channels to Alleviate Diabetes Mellitus via a Ca^2+^- CaMKKβ-AMPK pathway"

### Supplementary files

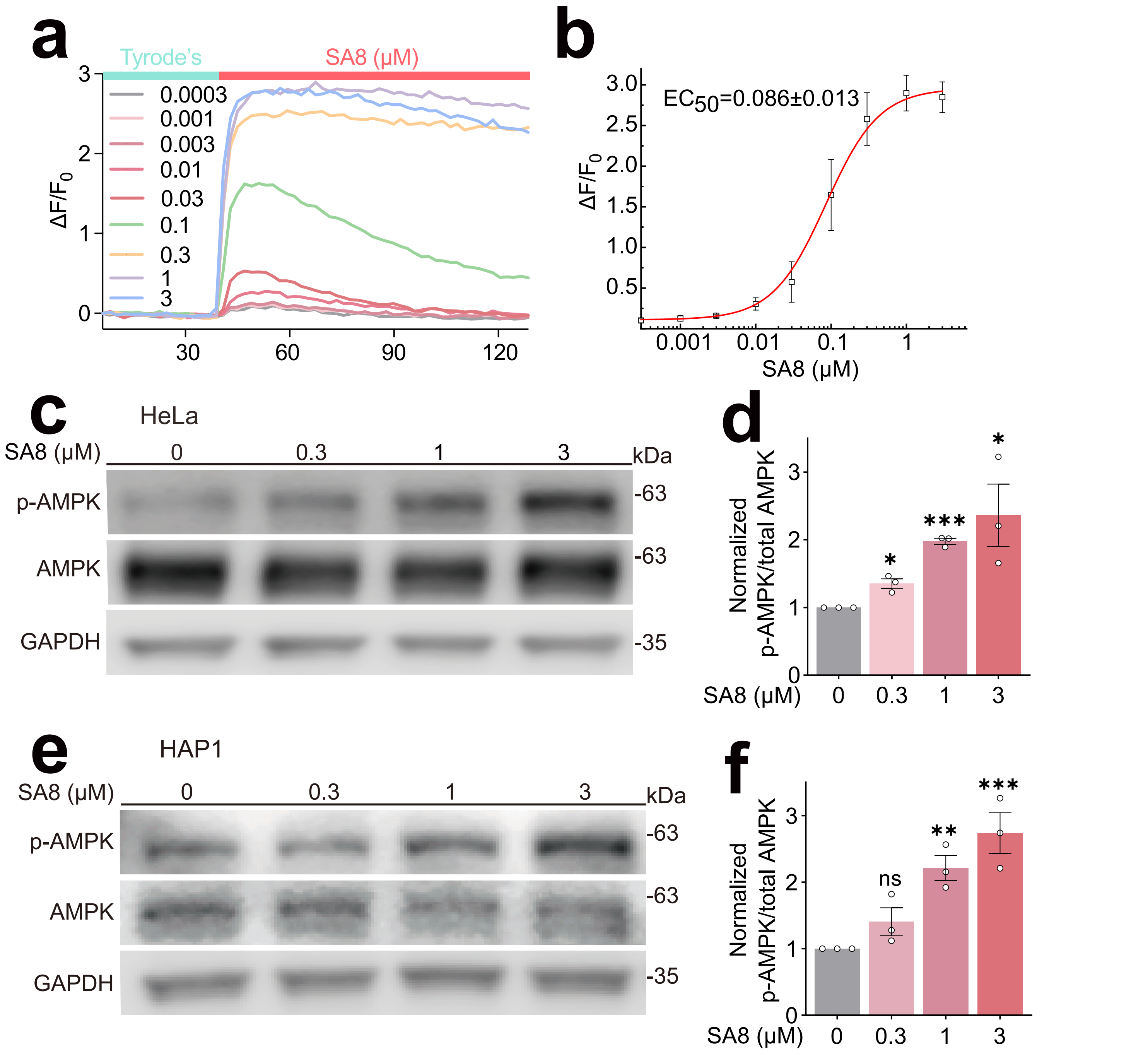

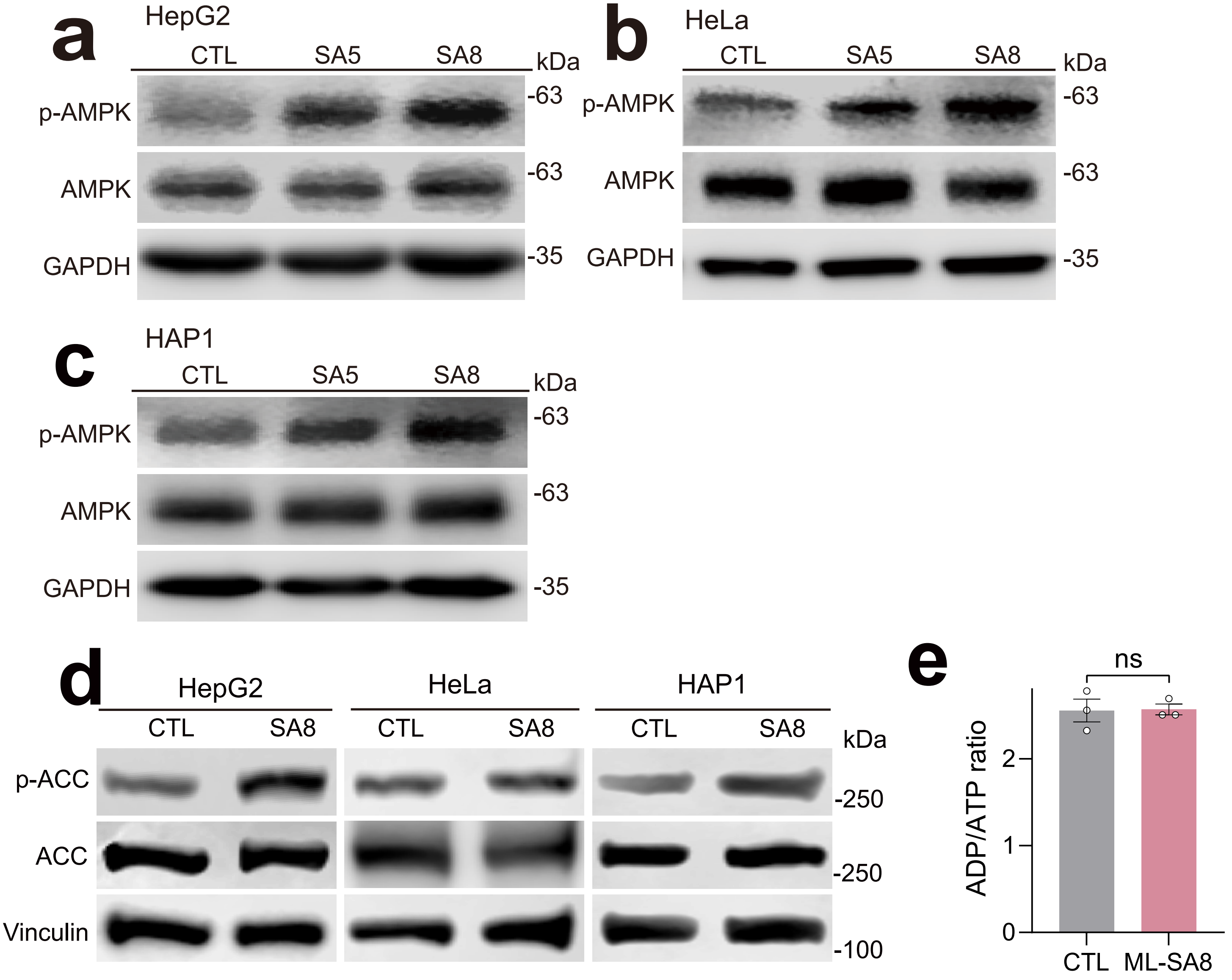


#### *Suppl. Fig.2*

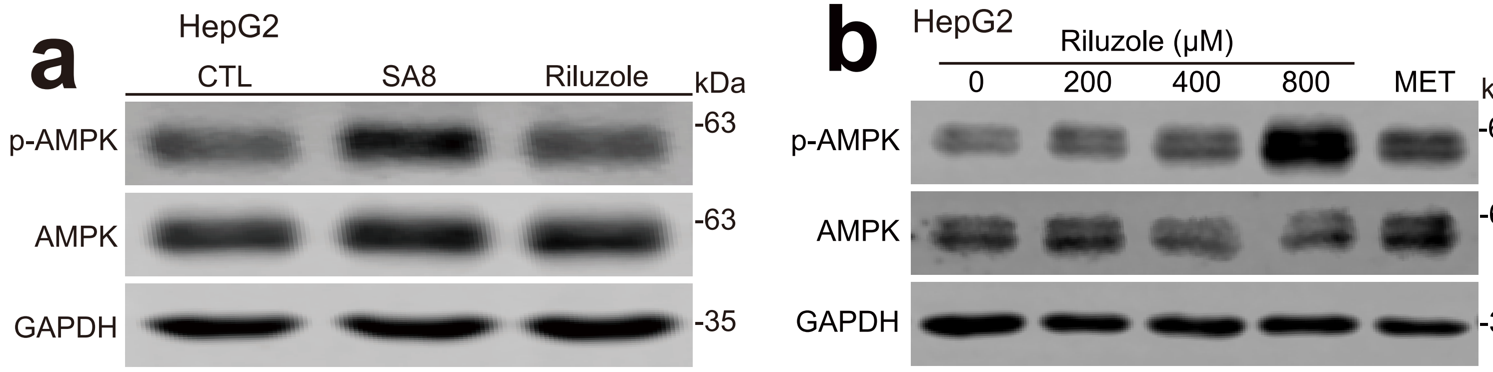


#### *Suppl. Fig.3*

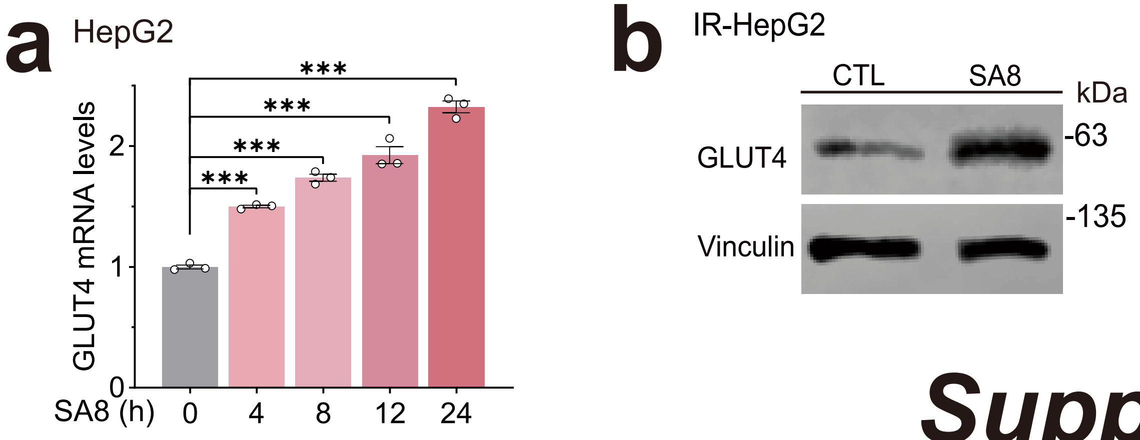


Suppl. Fig.4

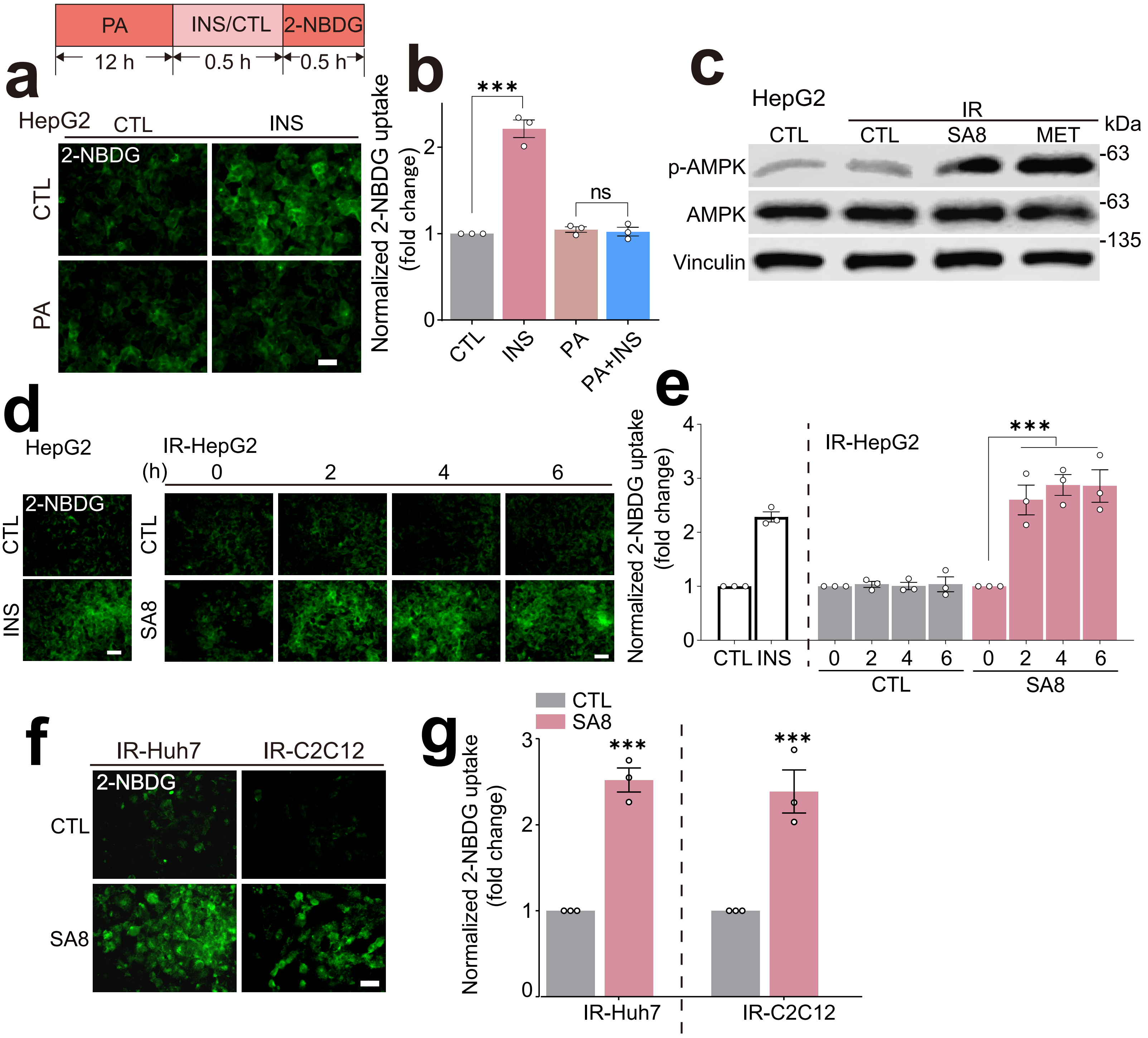


Suppl. Fig.5

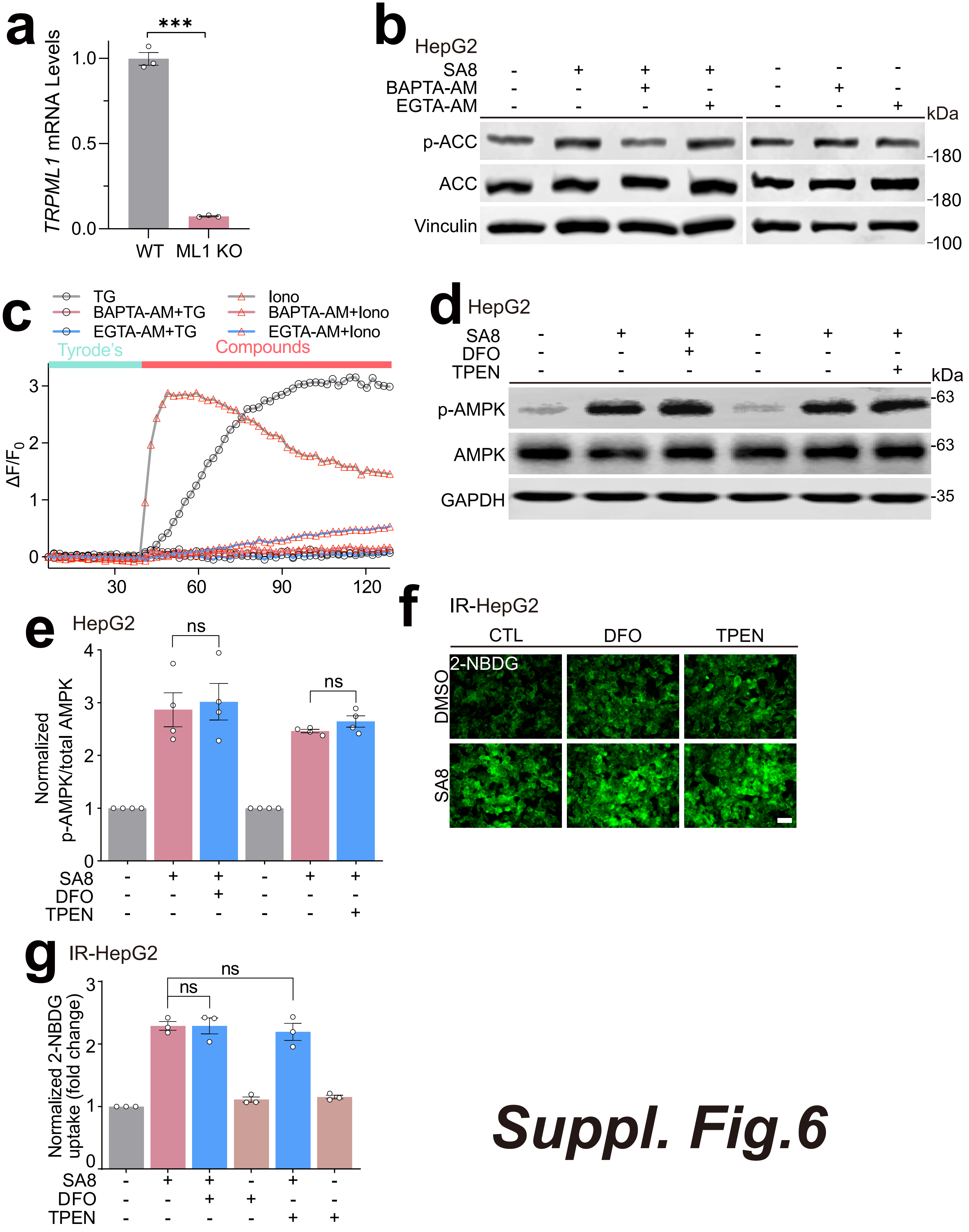

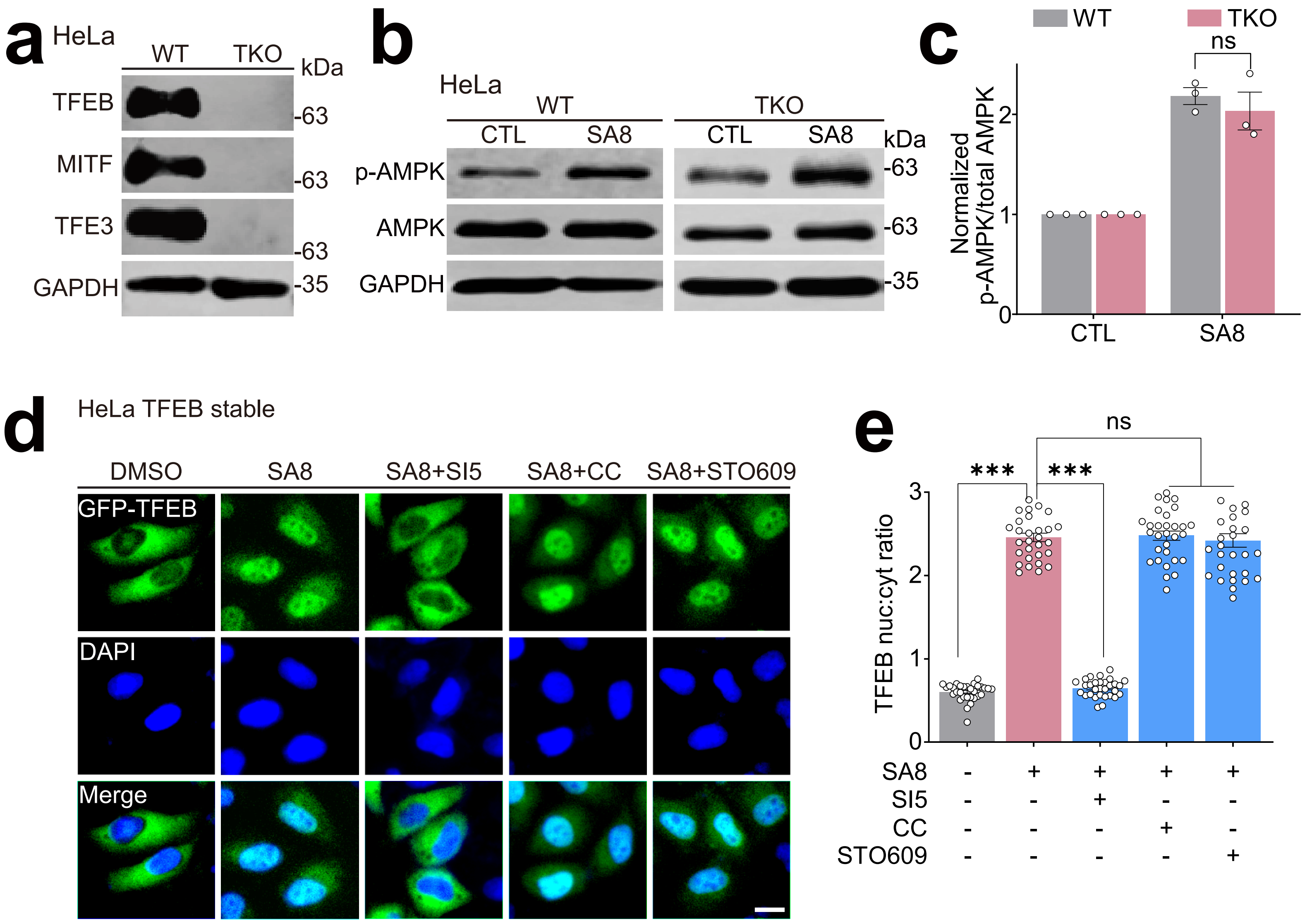

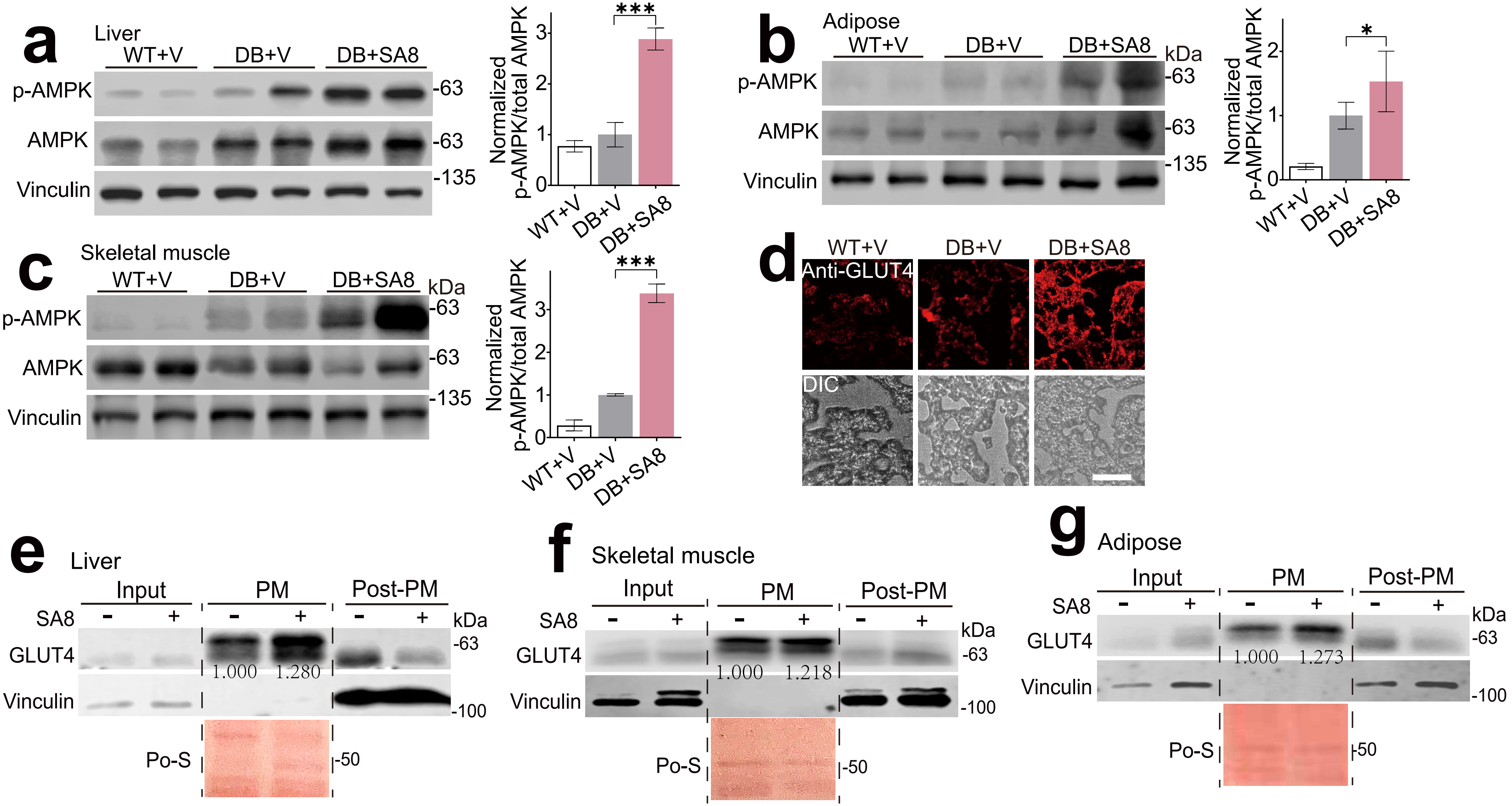

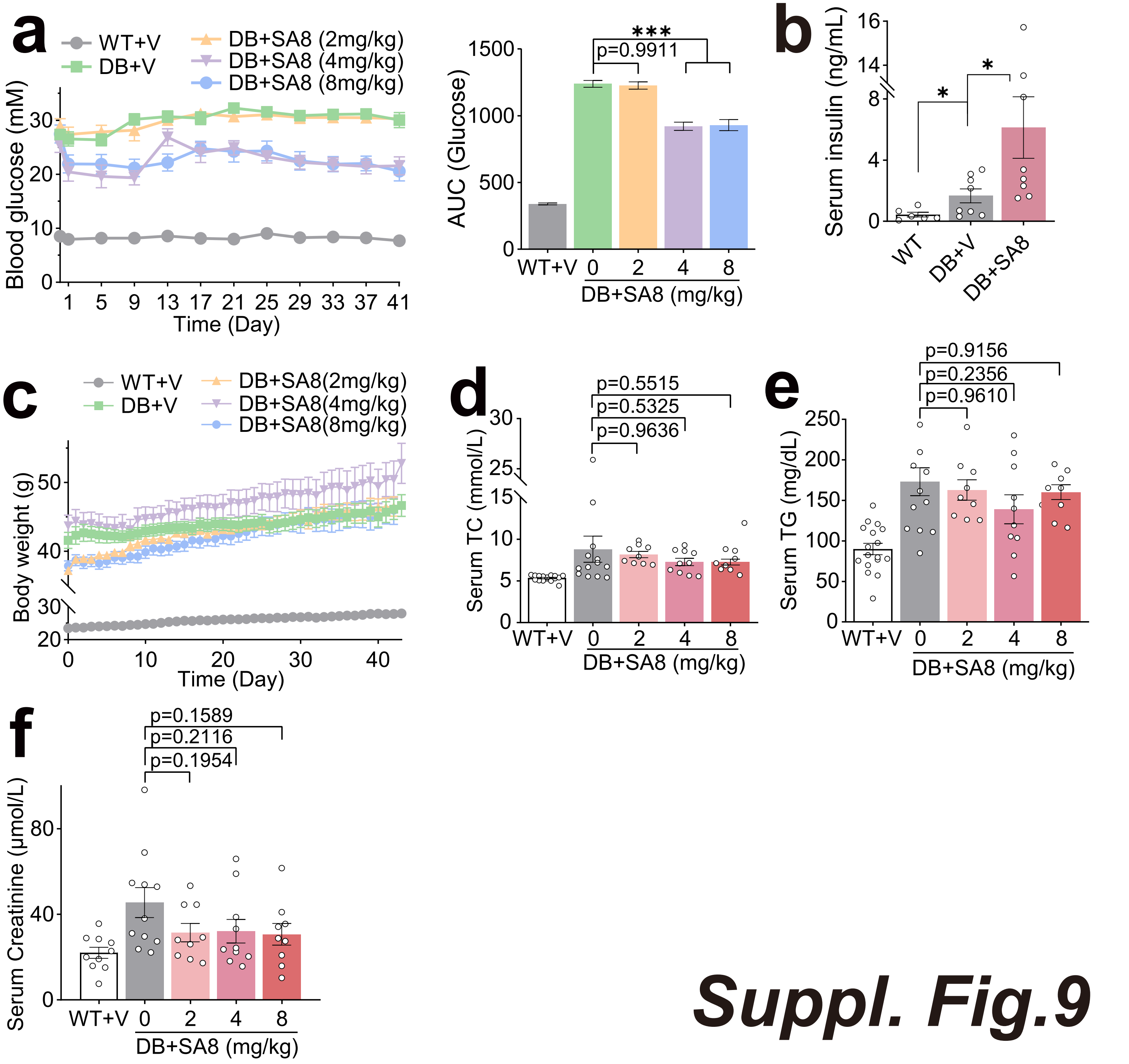

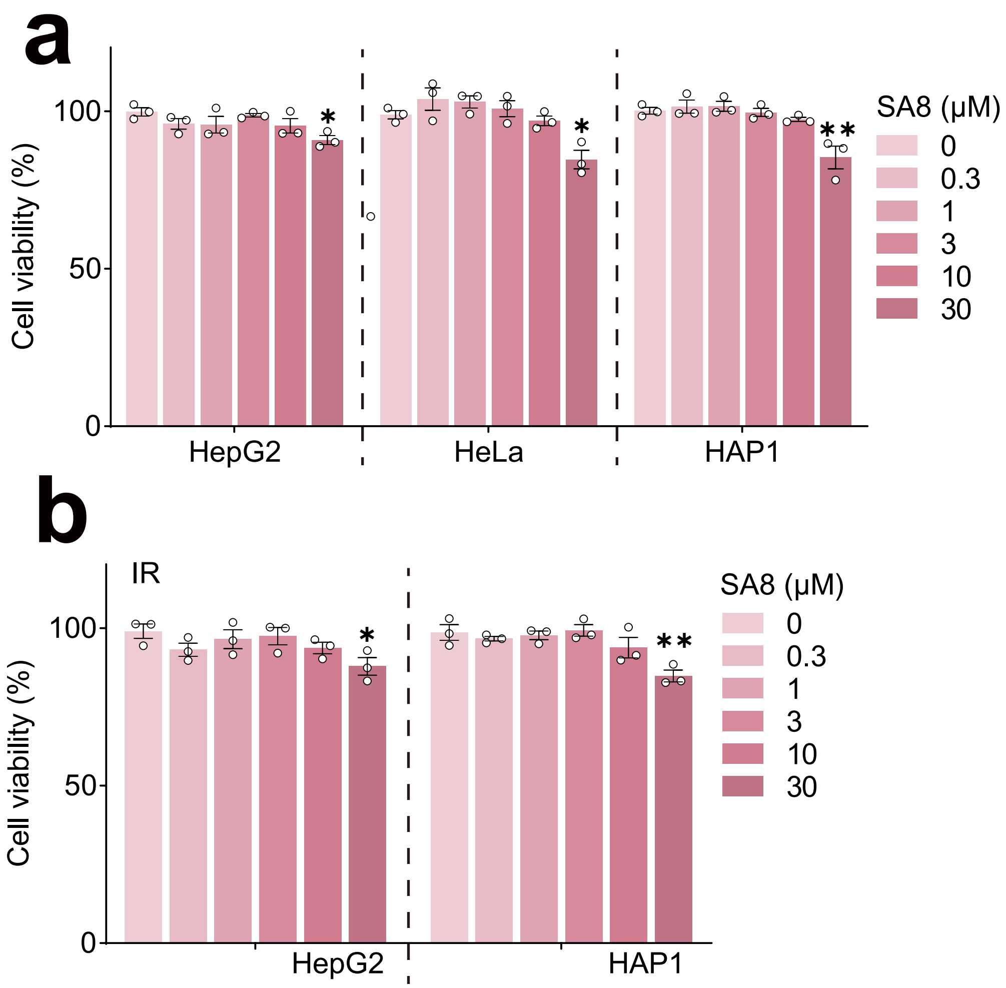
